# Theoretical causes of the Brazilian P.1 and P.2 lineages of the SARS-CoV-2 virus through molecular dynamics

**DOI:** 10.1101/2021.04.09.439181

**Authors:** Micael D. L. Oliveira, Kelson M. T. Oliveira, Jonathas N. Silva, Clarice Santos, João Bessa, Rosiane de Freitas

## Abstract

The new *β*-coronavirus has been causing sad losses around the world and the emergence of new variants has caused great concern. This pandemic is of a proportion not seen since the Spanish Flu in 1918. Thus, throughout this research, the B.1.1.28 lineage of the P.1 clade (K417T, N501Y, E484K) that emerged in Brazil was studied, as well as the latest Delta variant. This is because the molecular mechanisms by which phenotypic changes in transmissibility or mortality remain unknown. Through molecular dynamics simulations with the NAMD 3 algorithm in the 50*ns* interval, it was possible to understand the impact on structural stabilization on the interaction of the ACE2-RBD complex, as well as simulations in 30*ns* for the neutralizing antibody P2B-2F6, with this antibody was derived from immune cells from patients infected with SARS-CoV-2. Although not all molecular dynamics analyzes support the hypothesis of greater stability in the face of mutations, there was a predominance of low fluctuations. Thus, 3 (three) analyzes corroborate the hypothesis of greater ACE2-RBD stability as a result of P.1, among them: Low mean RMSF values, greater formation of hydrogen bonds and low solvent exposure measured by the SASA value. An inverse behavior occurs in the interaction with neutralizing antibodies, since the mutations induce greater instability and thus hinder the recognition of antibodies in neutralizing the Spike protein, where we noticed a smaller number of hydrogen bonds as a result of P.1. Through MM-PBSA energy decomposition, we found that Van der Waals interactions predominated and were more favorable when the structure has P.1 strain mutations. Therefore, we believe that greater stabilization of the ACE2-RBD complex may be a plausible explanation for why some mutations are converging in different strains, such as E484K and N501Y. The P.1 concern variant still makes the Spike protein recognizable by antibodies, and therefore, even if the vaccines’ efficacy can be diminished, there are no results in the literature that nullify them.

## 1. Introduction

In January 2021, Brazilian researchers identified a novel variant of the coronavirus, called P.1 (20J/501Y.V3), during a period of unprecedented increase in hospitalizations and incidence in cases of reinfection in the city of Manaus, where it was shown to be correlated with the emergence of novel lineage. From the genetic sequencing, it consists of multiple mutations, among the most worrying are 3 (three) that affected the RBD region of Spike glycoprotein: K417T, E484K and N501Y (Naveca et al., 2021). The P.1 strain identified has many similarities with the discovery in the United Kingdom B.1.1.7 (20H/501Y.V2 or VOC-202012/01). Since there has been an increase in transmissibility by 1.4-2.2 times, in addition to the greater probability of lethality by 1.1−1.8 times (Coutinho et al., 2021; Faria et al., 2021). According to data from the Epidemiological Surveillance System in Brazil in relation to the first three months of 2021, it was noticed that in the age groups from 30 to 39 years old there was an increase in cases by 565.08%, while in the age group from 40 to 49 years for a rate of 626%. Finally, when analyzing the death rate, the data are less expressive. Consequently, there is a shift towards younger age groups while the number of occurrences at older ages has been reduced (Castro, 2021).

The mutation rate for the SARS-CoV-2 virus is considered moderate and therefore lower than the rate for other viruses such as influenza and HIV. Possibly this result results from Nsp14, which has a correction and revision mechanism for mutations that eventually occur Duchene et al. (2020).

The P.2 lineage evolved independently in Rio de Janeiro without being directly related to the P.1 lineage (Voloch et al., 2021). We should note that the P.2 and South African lineage have the same mutations in common in the RBD region of the Spike protein. In South Africa, lineage B.1.351 emerged whose studies report greater difficulty in the recognition of antibodies (Moyo-Gwete et al., 2021; P. Wang et al., 2021). It is then constituted by eight mutations in the protein Spike, being 3 (three) of these: K417N, E484K and N501Y. It is noticeable the convergence of some mutations, specifically in position E484. Thus, several variants of the coronavirus, whether identified in South Africa, Brazil or the United Kingdom, carry this same mutation.

First of all, we have to highlight that theoretical and computational chemistry techniques are essential to see what is often inaccessible experimentally or even to provide new insights into the most intrinsic molecular mechanisms of diseases. Thus, molecular dynamics simulations are an essential tool for understanding the mechanisms involved in biomolecular interactions. Although, it is known that the precision of the results depends essentially on the elapsed time of simulation in addition to the precision of the adopted force field (Durrant & McCammon, 2011). In this research we studied the P.1 and P.2 variants impact using simulations of atomistic molecular dynamics (all-atom) for the ACE2-RBD and antibody-antigen complexes to find theoretical explanations of the causes of the increase in viral transmissibility or even severity of symptoms. In the end, we analyzed the RMSF atomic fluctuations, RMSD stability, formed Hydrogen bonds and SASA value of solvent exposure in the ACE2-RBD interaction. In addition, the non-covalent terms of molecular mechanics ⟨*E*_*MM*_⟩ were also estimated in addition to the total solvation energy by decomposing MM-PBSA energy.

## 2. Materials and methods

### 2.1. Mutagenesis in silico

As a result of the unavailability of crystallographic structures containing the mutations of the P.1 lineage (K417T, E484K and N501Y) and P2 (K417N, E484K and N501Y) that arose in Brazil, first it was necessary to use reference structures from the RCSB Protein Data Bank database (https://www.rcsb.org/) (Berman et al., 2000) where amino acid substitutions would be applied. The *in silico* mutation was performed in the PyMOL 2.3 software (Schrödinger, LLC, 2015) with the module “Mutagenesis” (see Figure 1) having the rotamer with lowest steric tension automatically defined by the software. In addition, the standard rotamer library was adopted, being dependent on the *ϕ* and *ψ* angles of the backbone. The prediction of new conformation of ACE2-RBD (PDB ID: 6M0J) (X. Wang et al., 2020) and antibody-antigen (PDB ID: 7BWJ) (Ju et al., 2020) with P.1 and P.2 mutations were performed with Branch-and-Prune implementation algorithm. After the insertion of mutations, it was necessary to remove steric clashes between the atoms and thus was performed molecular dynamics simulations to minimize and equilibrate the resulting structure.

**Figure 1.**
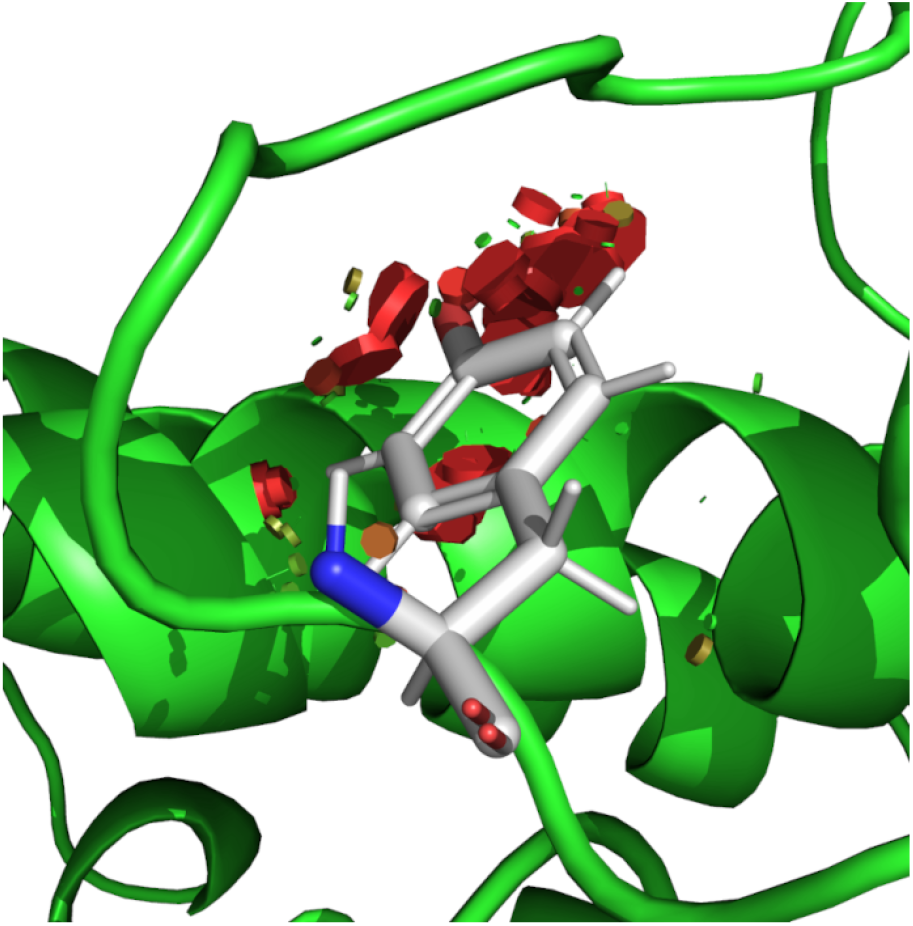
Insertion of the N501Y mutation is presented, belonging to the P.1/P.2 strains and applied to the ACE2-RBD complex (PDB ID: 6M0J). The software PyMOL (Schrödinger, LLC, 2015) was used for mutagenesis and presented a probability of tension at the best rotamer with a score of 31.18. The small green disks denote atoms that are almost in a steric or subtly overlapping conflict. On the other hand, the large red discs indicate a significant overlap of the Van der Waals radius.

In parallel, the I-Mutant 3.0 (Capriotti, Fariselli, & Casadio, 2005) tool was used, which implements the SVM machine-learning approach to predict ΔΔ*G* as a result of the emerging variants in the ACE2-Spike complex (PDB ID: 7DF4) and neutralizing antibodies RBD-Ty1 (PDB ID: 6ZXN). Thus, we adopt structures with a greater number of amino acids since the computational complexity is lower. We adopt the physiological condition at *pH* = 7.4, temperature of 25^*°*^*C* and SVM3 ternary classification algorithm where it denotes stability (ΔΔ*G ≥* +0.5*kcal · mol*^−1^), destabilization (ΔΔ*G* ≤ −0.5*kcal · mol*^−1^) and neutrality (−0.5*kcal · mol*^−1^ ≤ ΔΔ*G* ≤ +0.5*kcal · mol*^−1^). It is noted that although not all authors denote the positive sign as stabilization, it was agreed in this work that: + (Stabilization); − (destabilization) as shown in the equation below:

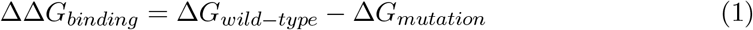

Finally, we also study the stability as a function of the mutations that constitute the P.1 and P.2 variants with the help of the Prime algorithm integrated into the Schrodinger Maestro 2020-3 Schrödinger, LLC (2020) software. Through the module “Residue Scanning” Beard, Cholleti, Pearlman, Sherman, and Loving (2013) the mutations of P.1 and P.2 were computationally inserted where the backbone was further minimized by the Prime algorithm in a cutoff of 5.0Å in relation to residues near the mutation area. A total of 11 mutations characterizing the P.1 variant (L18F, T20N, P26S, D138Y R190S, K417T, E484K, N501Y, D614G, H655Y, T1027I and V1176F) in the ACE2-Spike complex (PDB ID: 7DF4) reported were inserted be present in more than 75% of the sequenced samples. Finally, it was possible to decompose MM-GBSA energy by the Prime algorithm in the interaction between the ACE2 receptor and the Spike protein due to mutations, where we also calculated the values of Δ_*affinity*_, Δ_*stability*_, Δ_*energy*_. It is important to note that energy terms have two versions, that is, when analyzing stability and affinity. Therefore, in all mutations to adopt the same comparative criterion since not all of them appear in the ACE2-RBD interface, the energy analyzes corresponded to stability rather than affinity.

### 2.2. Parameters of molecular dynamics in NAMD

In this step, atomistic simulations (all-atom) of molecular dynamics were performed in the ACE2-RBD complex (PDB ID: 6M0J) (X. Wang et al., 2020) with an elapsed time of 50*ns* in front of the lineage P.1. We also studied the impact of P.1 on the interaction with specific neutralizing antibody P2B-2F6 with 30*ns* for RBD (PDB ID: 7BWJ) (Ju et al., 2020) that were derived from B lymphocytes from patients infected with SARS-CoV-2. All proteins were previously prepared with the software in its version for academic purposes Schrödinger Maestro 2020-3 using the “Protein Preparation” module, a step that included adjustments in ionization states, removal of water molecules, cofactors, assignment of partial charges and addition of Hydrogen atoms to the structure. The generation of all input and configuration files was done using the QwikMD 1.3 (Ribeiro et al., 2016) plugin implementing in the VMD 1.9.4.48a (Humphrey, Dalke, & Schulten, 1996) graphical interface. The complex was immersed in a solvation box (see Figure 3) with cubic geometry and applying PBC (periodic boundary conditions) containing water molecules described by the TIP3P model at a standard distance of 12.0 Å in relation to the frontiers of the box, as well as the addition of *Na*^+^ and *Cl*^−^ counterions to neutralize the system at a physiological molar concentration of 0.15 *mol · L*^−1^. In the ACE2-RBD complex, a total of 422, 710 atoms and 136, 320 water molecules were present, as determined automatically by the QwikMD algorithm. It is important to note that values can fluctuate slightly between the structure containing the mutations and the absence. In the antibody-antigen complex, 465, 723 atoms were present out of 151, 791 water molecules. Meanwhile, all topology files were generated with the CHARMM36 (Huang & MacKerell Jr, 2013) force field.

**Figure 2.**
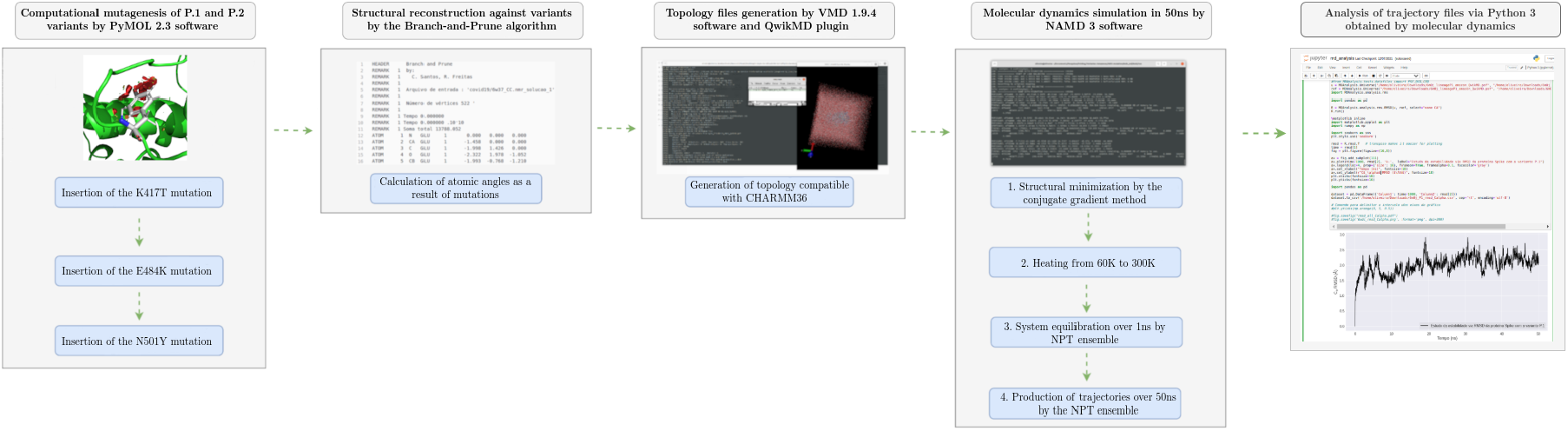
Steps taken from mutagenesis, structural minimization of the structures containing the mutations and proceeding to the final step of analysis for structural stability.

**Figure 3.**
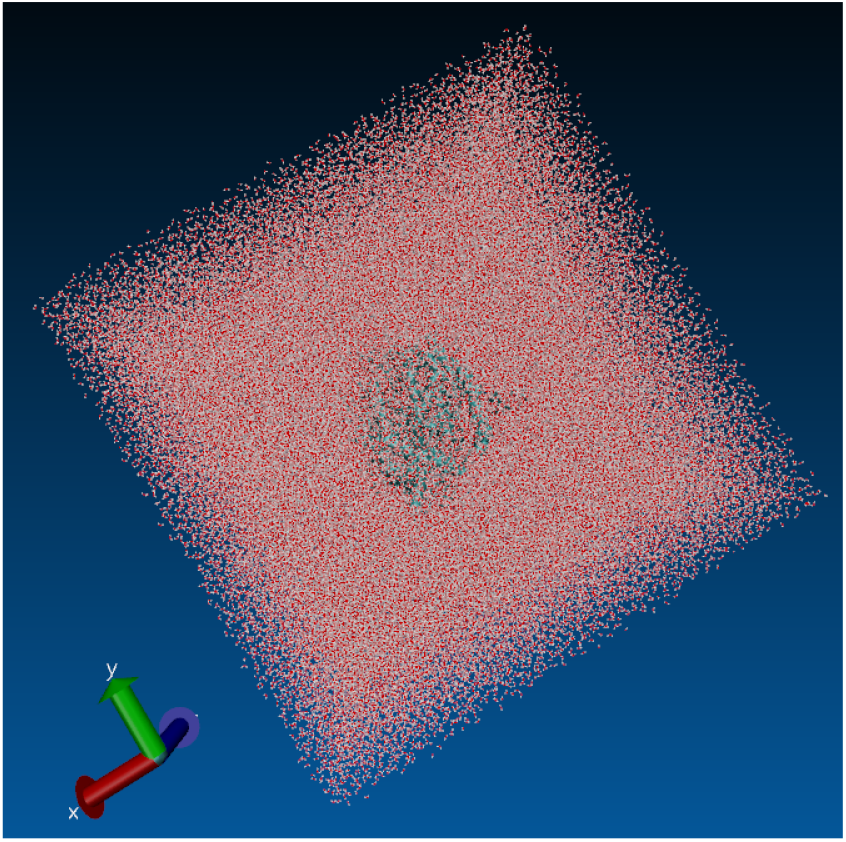
Cubic solvation box whose ACE2-RBD complex was immersed 12.0Å away from the boundaries with a volume of (163.57Å)^3^, and whose visualization was in the software VMD 1.9.4a48 (Humphrey et al., 1996).

The system was minimized using 1000 steps with conjugate gradient approach. Shortly thereafter, there was gradual heating of 60 − 300*K* at the rate of 1*K · ps*^−1^ under the NPT ensemble over 0.24*ns* and then the system was equilibrated in the NPT ensemble over 1*ns*. In addition, isotropic fluctuations were attributed to the simulation box, and for this the variable *FlexibleCell = no* was defined. Finally, the trajectory calculation step used the integration r-RESPA (Tuckerman, Berne, & Martyna, 1992) in an NPT thermodynamic cycle with the elapsed time of 8*ns* and recording every 20*ps*. Nosé-Hoover stochastic piston was used for pressure control similar to a barostat under the standard conditions of 1.01325*Bar*, period of 200*fs* and decay time with 100*fs*. While a Langevin dynamic thermostat was adopted with a coupling coefficient of *τ*_*p*_ = 1*ps*^−1^. In addition, long-range electrostatic interactions were estimated according to the Particle-Mesh-Ewald (PME) with a cutoff of 12.0angstrom formalism while the SHAKE (Kräutler, van Gunsteren, & Hünenberger, 2001) algorithm was used to constraint all hydrogens bonds along the simulation. Finally, the NAMD3 (Nanoscale Molecular Dynamics) (Phillips et al., 2020) algorithm was used to run all simulations with the acceleration of Nvidia GTX 1050 2 GB GPU with 640 CUDA cores at a time-step of 2*fs*. As a result of computational limitations, studies of mutations were restricted to the RBD region (229 amino acids) instead of the entire Spike receptor (1273 amino acids).

### 2.3. Parameters of molecular dynamics in GROMACS

In order to have a greater conviction of the results, we performed molecular dynamics simulations with the GROMACS 2019.1 software (Groningen Machine for Chemical Simulations) (Abraham et al., 2015). In this way, we simulate in the range of 50*ns* the recently published crystallography of the P.1 variant for the antigen-antibody containing mutations (PDB ID: 7NXB) (Dejnirattisai et al., 2021) and wild type (PDB ID: 7NX6) (Dejnirattisai et al., 2021). Meanwhile, the topology and coordinate files were generated using the force field CHARMM36 Huang and MacKerell Jr (2013). The system has been minimized by 5000 steps based on the steep descent method. Soon after, the system was heated to 300*K* until it reached stability with a convergence of *<* 1000*kJ · mol*^−1^ *· nm*^−1^. Then the equilibrium was performed for 125*ps* with an integration time of 1*fs*, with the canonical ensemble NVT at the elapsed time of 1*ns*. All native structures were immersed in a cubic solvation cell, using the TIP3P water model (Jorgensen, Chandrasekhar, Madura, Impey, & Klein, 1983), whose distance from the complex to the box boundaries was 10.0Å. Finally, the production of trajectories was performed using the NPT canonical cycle. It is noteworthy that the simulation box was infinitely replicated in a three-dimensional space under periodic boundary conditions. The time elapsed in each simulation was 50*ns* with trajectories recorded every 50*ps*. Long-range electrostatic interactions were estimated using the Particle-Mesh Ewald (PME) formalism (Darden, York, & Pedersen, 1993). Ions of *Na*^+^ and *Cl*^−^ were inserted to neutralize the system at the physiological concentration of 0.15*M*. The Parrinello–Rahman barostat (Parrinello & Rahman, 1980) was used to keep the pressure constant at 1*atm*, while the Berendsen algorithm (Berendsen, Postma, van Gunsteren, DiNola, & Haak, 1984) acted as a thermostat to keep the system temperature constant at 300*K*. The angles of the covalent bonds were constrained with the LINCS36 (Linear Constraint Solver) algorithm (Hess, Bekker, Berendsen, & Fraaije, 1997) and the geometry of the water molecules was adjusted using the SETTLE37 algorithm (Miyamoto & Kollman, 1992).

### 2.4. Molecular dynamics analysis

In the end, the conformational changes in the macromolecule as a result of the mutations were measured, through the temporal evolution RMSD and RMSF of the atomic displacement in relation to the *Cα* of the ACE2-RBD complex where frame 0 was adopted as a reference. The analysis of trajectories and construction of RMSD and RMSF graphics were possible with the MDAnalysis (Michaud-Agrawal, Denning, Woolf, & Beckstein, 2011) and Matplotlib libraries implemented in the Python 3 programming language. The fraction of native contacts was calculated based on the first frame of the MD simulations as a native conformation. Throughout the analysis of native contacts, the MDAnalysis tool where the function *radius_cut*_*q*_(*r, r*0, *radius*) was adopted, with the cut radius *<* 4.5Å. The calculation of the hydrogen bonds formed during the simulation was made using a plugin present in the VMD 1.9.4a.51 software where the donor-acceptor cut distance was 3.0Å while the cut angle was of 20^*°*^. Throughout the identification of saline bridges, a plugin was also used in this same software with an oxygen-nitrogen cutoff distance of 3.2Å. The estimate of the SASA value refers to the average of the frames, and the algorithm used was a plugin in the VMD with a cut radius of 1.4Å. Regarding the impact of P.1 and P.2 strains on the recognition of antibodies, one complex antigen-antibody was subjected to molecular dynamics simulations in the range of 30*ns* and later the mean values of RMSD, RMSF, SASA, in addition to formed hydrogen bonds.

The estimation of the error in the RMSD, RMSF, SASA and Hydrogen bonds values were based on the standard deviation of the measurements over time. We used the Scipy library in Python for all statistical analysis with the command *ttest_ind*(*a, b, equal_var* = *False*), where (*a, b*) denote two matrices containing the molecular dynamics RMS data. We analyzed the statistical significance of the comparative results using the two-way Student t-test to confirm the hypothesis of the difference between two means (*µ*_1_ *≠ µ*_2_) of the RMSD and RMSF values obtained from a set of statistically independent data and variance 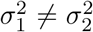. In order to justify the validity of the Student’s t-test approach, the Shapiro-Wilk normality test was performed for the number of hydrogen bonds, with a minimum confidence interval of 87%. Levene’s test was applied to verify equality between variances where *p >* 0.05, and therefore the inequality had to be considered throughout the hypothesis test. Consequently, we believe that the proposal of the t-test formalism for comparative analysis of MD simulations is justified. We considered the value *p <* 0.05 for a statistically significant difference for a confidence interval of 95%.

### 2.5. MM-PBSA energy decomposition

The decomposition of the energy values ⟨*E*_*MM*_⟩, ⟨ *E*_*elec*_⟩, and ⟨*E*_*V dW*_⟩ refer to the average of all molecular dynamics frames. All components of molecular mechanics and solvation terms were estimated for the ACE2-RBD complex (PDB ID: 6M0J) in the presence and absence of the P.1 strain, in which the explicit water molecules and ions were previously removed for analysis. Furthermore, throughout the analyzes for the antibody-antigen complex Spike S1 and P2B-2F6 we considered only the interaction of Spike S1 with the antibody light (L) chain, as a result of presenting more consistent potential energy results. We used the NAMDEnergy plugin to calculate *E*_*MM*_ while the term ⟨*SASA*⟩ was previously obtained. From the average solvation area it was possible to calculate the term 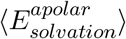 where the empirical values *γ ≈* 0.00542 *kcal · mol*^−1^ *·* Å^−2^ and *β ≈* 0.92*kcal · mol*^−1^ were adopted. The 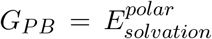 component of Poisson-Boltzmann was estimated for the last frame by the finite difference method using the DelPhi v8.4.5 algorithm (Li et al., 2012) with a 0.0001 convergence criterion, in a 0.10*M* solution having a dielectric constant for the solute with *ϵ* = 2.0 and for the solvent the water dielectric constant was *ϵ* = 80.0 water probe radius of 1.4Å was considered. It is important to note that the polar solvation term was considered implicit continuum solvation by the DelPhi algorithm. As a result of the high computational complexity and the inherent uncertainty in the estimation of the entropic term *T ·* Δ*S*, we did not include the MM-PB(SA) decomposition of this research.

The equation to estimate the polar contribution corresponds to the finite difference of the solution of the Poisson-Boltzmann equation (PB) (Homeyer & Gohlke, 2012; C. Wang, Greene, Xiao, Qi, & Luo, 2018):

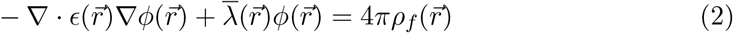

The entropic term is generally estimated only by normal or quasi-harmonic analysis and is the most complex component to obtain. The nonpolar solvation free energy stems from Van der Waals interactions between the solute and the solvent. The value of *SASA* denotes the surface area accessible to the solvent Genheden and Ryde (2015):

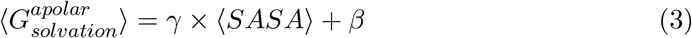

The solvation free energy is then estimated by the sum of Δ_*PB*_ associated with the Poisson-Boltzmann term and by the term 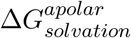, making up respectively the polar and nonpolar contributions to the solvation free energies. The polar contribution is typically obtained by solving the PB equation, while the term apolar is estimated from a linear relationship with the surface area accessible to the solvent (SASA) (Genheden & Ryde, 2015).

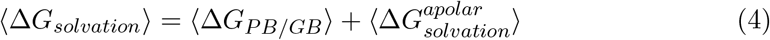

The term *E*_*MM*_ corresponds to the total molecular mechanics (MM) energy referring to the sum of the covalent interactions *E*_*int*_ with the non-covalent *E*_*V dW*_ and *E*_*elec*_. The internal energy term *E*_*int*_ consists of the sum of the energies associated with the chemical bonds *E*_*bond*_, the angles *E*_*angles*_ and the dihedral torsions *E*_*torsions*_ (Genheden & Ryde, 2015).

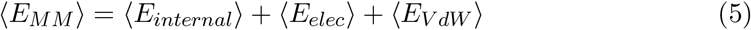

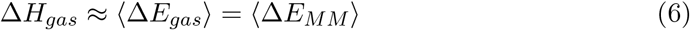

In the MM-PBSA method, the free energy of binding results from the sum of the interaction energy of the complex associated with the formation in its gaseous phase (vacuum), the term associated with total solvation and finally the entropic penalty:

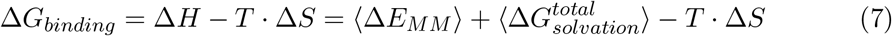

## 3. Results and Discussions

### 3.1. Study of the Brazilian lineage B.1.1.28 of clades P.1 and P.2

First of all, the structural reconstruction of the Spike protein containing the P.1 and P.2 mutations was possible by implementing an implicit enumeration algorithm called Branch-and-Prune (BP). It is therefore a method that lists all possible atoms and positions while discarding invalid ones. After mutagenesis and protein reconstruction, it was finally possible to perform the various analyzes of chemical interactions, molecular dynamics and MM-PBSA decomposition.

#### 3.1.1. *Stability analysis* ΔΔ*G*

Through an analysis with the aid of the I-Mutant 3.0 tool (Capriotti et al., 2005), the global value of ΔΔ*G* in the ACE2-RBD structure (PDB ID: 7DF4) as a result of the 3 (three) mutations that constitute the Amazonian lineage B.1.1.28/P.1 was −1.85*kcal · mol*^−1^ where: N501Y (−0.08*kcal · mol*^−1^); K417T (−1.50*kcal · mol*^−1^); E484K (−0.27*kcal · mol*^−1^). Thus we can see that the N501Y mutation that appeared in the UK a priori would not significantly affect the ACE2-RBD interaction. The same conclusion refers to the E484K mutation in South Africa that did not reach the critical threshold of −0.5*kcal · mol*^−1^. On the other hand, K417T that emerged in Amazonas generated an unexpected destabilization in the ACE2-RBD recognition, which although it can decrease the probability of cell invasion could make it difficult to recognize neutralizing antibodies induced by vaccines. In addition, the critical destabilization due to K417T could have a correlation with the unprecedented increase in January 2021 in infections and deaths in Amazonas due to COVID-19. The P.2 clade lineage is distinguished by the K417N mutation that generates less significant destabilization although still critical of −1.35*kcal · mol*^−1^. It is noted that mutations where there is destabilization tend to disappear due to evolutionary pressure since they cause a negative impact on the protein function (Ancien, Pucci, Godfroid, & Rooman, 2018). We must note that these instability results are only predictions, and only with molecular dynamics can we be more convinced of the conclusions. As reported in the literature, the phenomenon of stability is proportional to a lower conformational entropy, but it is not possible to infer that greater stability is necessarily related to the greater severity of the disease.

In order to have greater confidence in the stability results of the ACE2-RBD complex, we predicted ΔΔ*G* using the mCSM (Pires, Ascher, & Blundell, 2013) tool where: N501Y (−0.757*kcal · mol*^−1^); K417T (−1.594 *kcal · mol*^−1^); E484K (−0.081 *kcal · mol*^−1^). Again, the K417T mutation of the P.1 lineage showed a critical destabilization, as occurred in the I-Mutant 3.0 tool. When the P.2 lineage was analyzed, the K417N mutation generated a critical destabilization of −1.582 *kcal · mol*^−1^ providing preliminary evidence of changes in residue Lys417 in the Spike protein may be the main responsible for the increase in the transmissibility of the virus in the Amazonas. Through the tool DynaMut2 (Rodrigues, Pires, & Ascher, 2021) it was possible to understand the impact of mutations on chemical bonds (see Figure 4). The K417N mutation belonging to the P.2 lineage is considered destabilizing as a result of the loss of 1 (one) Hydrogen bond, 3 (three) hydrophobic contacts in addition to 2 (two) polar interactions at the ACE2-RBD interface. The same pattern was observed in the P.1 lineage with the K417T mutation, although with greater stability as a result of having lost fewer interactions with the disappearance of 1 (one) hydrogen bond, 1 (one) hydrophobic contact and 1 (one) interaction polar.

**Figure 4.**
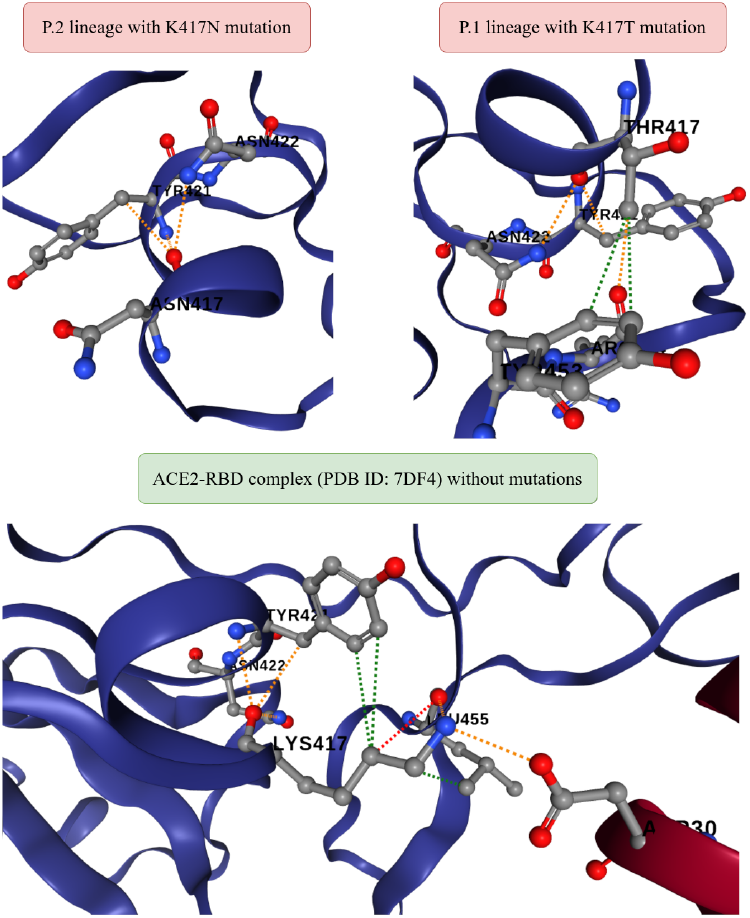
Comparison between the chemical interactions formed in the ACE2-RBD complex (PDB ID: 6M0J) as a result of the K417T and K417N mutations related to the P.1 and P.2 lineages, respectively. All diagrams were generated on the DynaMut2 (Rodrigues et al., 2021) platform. The dashed lines in green represent the hydrophobic contacts, the lines in red correspond to the Hydrogen bonds and finally the orange color refers to the polar interactions.

In order to deepen the analysis of ΔΔ*G* as a result of mutations (see Figure 3.1.1), we used public data obtained from the COVID-3D platform (http://biosig.unimelb.edu).au/covid3d/). As a result of the analyzes being to Spike’s trimeric complex, we restricted the results to the mean values of ΔΔ*G* among the 3 (three) chains. Even though there is no total consensus among the total of 5 (five) tools used, there is a predominance of structural destabilization as a result of K417T where the average value was 0.441*kcal · mol*^−1^. Similarly, the K417N mutation showed a destabilization with a mean value of 0.447*kcal · mol*^−1^. Based on these results, the two mutations possibly have similar phenotypic behavior in the virus, even if they belong to different strains.

We studied the stability as a function of P.1 and P.2 mutations with the help of the Prime algorithm belonging to the Schrodinger Maestro 2020-3 software (Schrödinger, LLC, 2020). By means of the module “Residue Scanning” (Beard et al., 2013) the mutations of P.1 and P.2 were computationally inserted where the backbone was later minimized by the Prime algorithm. In this way, the decomposition of MM-GBSA energy was possible. According to the results presented in the Table 2, we can see that the N501Y mutation is the one that most favors the structural stability in the ACE2-RBD interaction. Consequently, according to the data obtained, it would be the most worrying mutation and one that would be more correlated with changes in transmissibility and lethality. Unexpectedly, the K417T mutation was not critical compared to other mutations already existing in other strains of the new coronavirus. This conclusion is visible in all energy contribution terms, whether Δ*E*_*V dW*_, Δ*G*_*solvation*_ or Δ*E*_*elec*_.

**Table 1.** Results obtained by the Prime algorithm when analyzing the impact of 11 (eleven) mutations that affected the Spike protein as a result of the P.1 variant. These analyzes were obtained from the ACE2-Spike trimeric complex (PDB ID: 7DF4). The notation Δ corresponds to a comparison between the results obtained with the mutation in relation to its absence. The energy impacts (*kcal · mol*^−1^) correspond to the interaction between the A chain (ACE2 receptor) and the 3 (three) chains of the Spike protein.

| Mutation | $\Delta$ Affinity | $\Delta$ Stability | $\Delta$ Energy | $\Delta E_{VDW}$ | $\Delta E_{elec}$ | $\Delta G_{solvation}^{GB}$ | $\Delta SASA$ |
| --- | --- | --- | --- | --- | --- | --- | --- |
| L18F | 0.000 | +16.348 | -7.412 | +11.698 | +12.089 | -14.915 | -37.201 |
| T20N | 0.000 | +13.099 | -102.641 | +7.955 | -114.172 | -7.910 | +111.414 |
| P26S | 0.000 | +5.267 | -91.843 | +0.305 | -59.103 | -3.967 | -4.532 |
| D138Y | 0.000 | +2.039 | +84.209 | +16.434 | +176.809 | -111.311 | -29.831 |
| R190S | 0.000 | +74.425 | +149.275 | +26.718 | -36.009 | +166.855 | +173.849 |
| K417T | +2.943 | +2.239 | +22.219 | +11.207 | -53.042 | +98.945 | +84.866 |
| E484K | +2.175 | +32.268 | +77.898 | +3.583 | +255.865 | -199.598 | +194.320 |
| N501Y | -7.969 | +71.708 | +175.868 | +8.243 | +99.376 | +5.815 | -13.546 |
| D614G | 0.000 | -5.740 | +97.340 | +5.902 | +6.811 | +69.985 | +187.280 |
| H655Y | 0.000 | -16.133 | -18.953 | +5.049 | -3.283 | +0.118 | -96.247 |
| T1027I | 0.000 | -13.365 | +39.435 | -7.604 | +13.057 | +14.826 | -85.637 |

**Table 2.** MM-GBSA energy analyzes referring to the antibody-antigen complex (PDB ID: 6XEY).

| Mutation | $\Delta$ Affinity | $\Delta$ Stability | $\Delta$ Energy | $\Delta E_{VDW}$ | $\Delta E_{elec}$ | $\Delta G_{solvation}^{GB}$ | $\Delta SASA_{total}$ |
| --- | --- | --- | --- | --- | --- | --- | --- |
| D138Y | +0.103 | +105.634 | +187.810 | +27.800 | -31.749 | +77.766 | -190.254 |
| R190S | +0.001 | +54.771 | +129.620 | +22.340 | -45.077 | +155.585 | +272.830 |
| K417T | -2.597 | -81.126 | -61.140 | +7.130 | -52.061 | +75.545 | +9.647 |
| E484K | +11.136 | +27.322 | +72.950 | +6.790 | +166.127 | -122.500 | -11.093 |
| N501Y | -0.017 | +64.433 | +168.590 | +3.230 | +61.129 | +22.341 | -102.301 |
| D614G | -0.001 | -0.001 | +100.300 | +18.300 | -23.721 | +96.411 | +69.057 |
| H655Y | 0.000 | +145.107 | +19.050 | +150.550 | -69.125 | +3.477 | -149.632 |
| T1027I | 0.000 | +6.080 | +58.870 | -2.590 | +38.942 | +3.105 | -77.742 |

It should be noted that the stability prediction ΔΔ*G* in different tools indicates different conclusions. However, there is a predominance of structural stability against mutations, which will be corroborated by molecular dynamics.

Using the PDBePISA Krissinel (2010) platform, the variant P.1 presented Δ_*i*_*G ≈* −6.5 *kcal · mol*^−1^, and therefore more favorable compared to the structure without any mutations where Δ_*i*_*G ≈* −5.9 *kcal · mol*^−1^. This would perhaps explain *a priori* why the P.1 line would be characterized by an increase in transmissibility Faria et al. (2021).

When analyzing the RBD-Ty1 interaction (PDB ID: 6ZXN) with the mutations present in the P.1 strain, a destabilizing behavior was found, which reflects a less interaction between the Spike glycoprotein and monoclonal antibodies so that: N501Y (0.35*kcal · mol*^−1^); K417T (−0.90*kcal · mol*^−1^); E484K (−0.50*kcal · mol*^−1^). Among all the results, the most worrying is precisely the K417T mutation that emerged in South Africa, which presents a critical destabilization and may a priori hamper the immune response through the administration of vaccines. Regarding the P.2 strain, the K417N mutation showed lower stability compared to the K417T, resulting in −0.82*kcal · mol*^−1^. Lastly, when we performed the protein-protein docking (see Table 3.1.1) on the HDock platform (http://hdock.phys.hust.edu.cn/) (Yan, Zhang, Zhou, Li, & Huang, 2017) with all default settings from the last frame of MD simulations, where the ACE2-RBD complex (PDB ID: 6M0J) containing the lineage P.2 resulted in a greater affinity with a relative score of −311.75 while in the absence of mutations it was −310.19. In highlight we have the P.1 variant, which presented an even more expressive interaction with −334.69. These results corroborate the hypothesis that P.1 makes Spike interaction with the ACE2 cell receptor more expressive. The PatchDock Schneidman-Duhovny, Inbar, Nussinov, and Wolfson (2005) platform was also used, which employs principles of geometric complementarity where we adopt the “Default” configuration and the type of protein-protein complex in all simulations and RMSD clustering of 4Å. Thus, P.2 mutations also reflected an increase in affinity in the ACE2-RBD complex where the score increased from 15722 to 16940 compared to the reference structure. When analyzing all the mutations that constitute P.1/P.2, we realized that the N501Y mutation was the only one that directly affected the antibody-antigen interface although the value of ΔΔ*G* was not significant, because otherwise, the harm to patients could be even greater than it currently is.

### 3.2. Quantum chemistry modelling

From the analysis of chemical-quantum descriptors (see subsection 3.2), we can see that the transition from Lysine (K417) to Threonine (417T) provides greater electron affinity, ionization potential and electronegativity. All these are factors that can contribute to greater interaction in the ACE2-RBD interface.

#### 3.2.1. Impact of P.1/P.2 through MD analysis

It should be noted that the mutations that constitute the South African strain affected the same positions in the RBD region of the Spike protein in relation to the Brazilian strain B.1.1.28/P.1. Thus, the simulations presented here will present similar conclusions for both strains. Although molecular dynamics is a rigorous approach to study the interaction of biomolecules, it is necessary to calculate as many trajectories as possible to minimize statistical errors. It is very difficult to analyze seemingly random behavior. At first glance, it seems impossible since there is no visible pattern. In molecular dynamics, when there is a larger sample, the behavior is not completely random, revealing intrinsic patterns of the analyzed system. Thus although the results presented here have a relatively short time of 50*ns* for ACE2-RBD and 30*ns* for RBD + P2B-2F6 antibody, we believe that the results are consistent and relevant. Despite the widespread use of MD simulations, little statistical approach is used to attest to the significance of the results to minimize subjectivity in the conclusions. This may be due mainly to the low sampling of data resulting from molecular dynamics, which tends to make statistically significant even very small variations. Despite of all, a statistical approach is found in the literature that used ANOVA to study the impacts on the activation of the GPCR protein (Bruzzese, Dalton, & Giraldo, 2020). It should be noted that the tests of statistical significance only make sense when running simulations at a time adequate to the studied problem. And since in this case we are only addressing the stability in the protein-protein interaction, we should have a time greater than 50*ns* − 100*ns* for valid results, whether from the physical or statistical aspect. Furthermore, as a consequence of the thermodynamic non-equilibrium state characteristic of biological systems, we used Langevin’s dynamics to obtain more consistent results. For this reason, we used the Student’s t-test to confirm whether the differences in the conformational fluctuations of the P.1 strain are really significant. Our criterion for quantifying the influence of structural flexibility in the presence and absence of the P.1 strain was the value of RMSD and also for the analysis as a function of amino acids by RMSF. This difference corresponds to the variation of the average values RMSD and RMSF between the structure with the mutations of P.1 and the wild-type structure, which was important for the tests of statistical significance (see *Supplementary Material*).

By constructing a 2D graph (see Figure 8) for the interactions between residues from the last frame of molecular dynamics, we can better visualize the impact of mutations on the ACE2-RBD and antibody-antigen interaction. Therefore, it was noted that in the face of variant P.1, 3 (three) bonds of the cation-*π* type (in red) were formed in addition to 1 (one) *π*-stacking bond (in dark blue). It is important to note that although interactions tend to change along the molecular dynamics, we chose the last frame for more consistent results.

**Figure 5.**
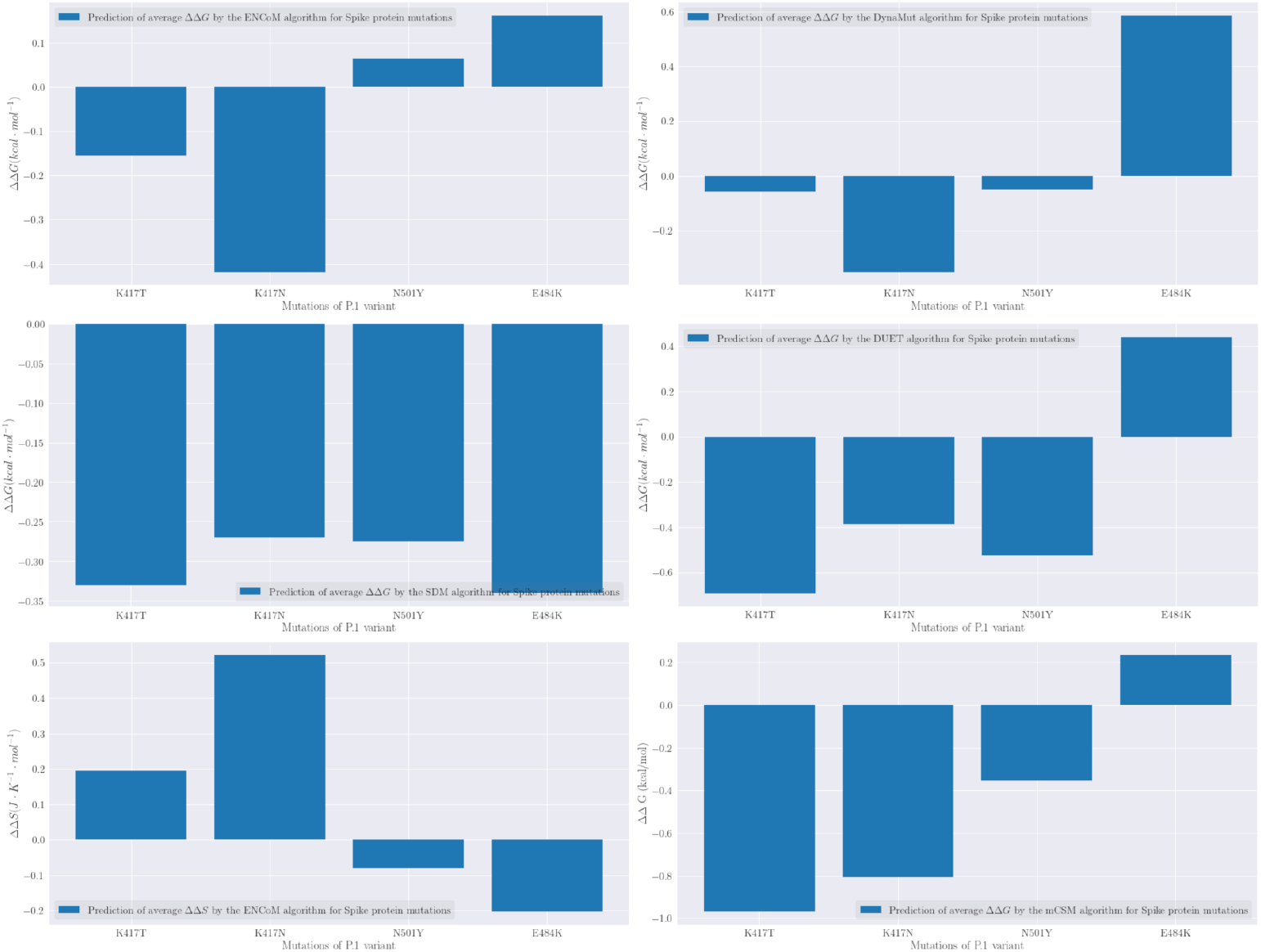
Prediction of the values of ΔΔ*G* performed by 5 (five) machine-learning algorithms: mCSM, EnCoM, DUET, SDM, DynaMut. Thus, we analyzed the mutations that constitute the P.1 strain and that affected the Spike glycoprotein in complex with ACE2-B0AT1. All data were extracted from the COVID-3D database Portelli et al. (2021).

**Figure 6.**
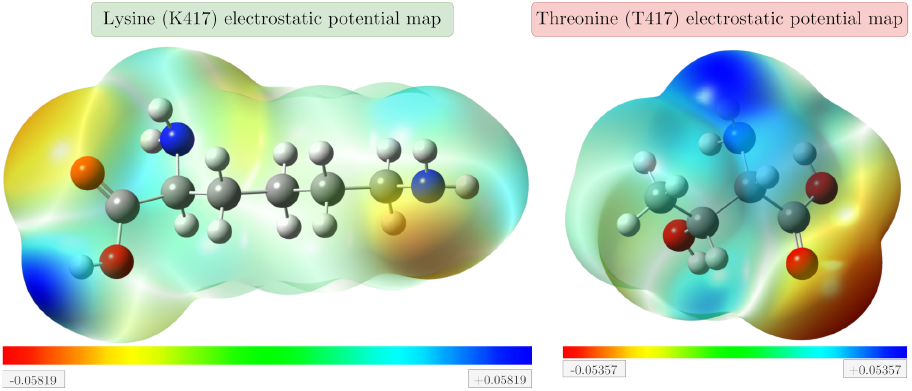
Electrostatic potential maps (MEPS) of amino acids affected by the P.1 variant, which were optimized at the DFT theory level with the Gaussian bases 6-311G (2d, 2p) with the hybrid functional B3LYP in the Gaussian 09W software. All figures were generated by GaussView 6 software.

**Figure 7.**
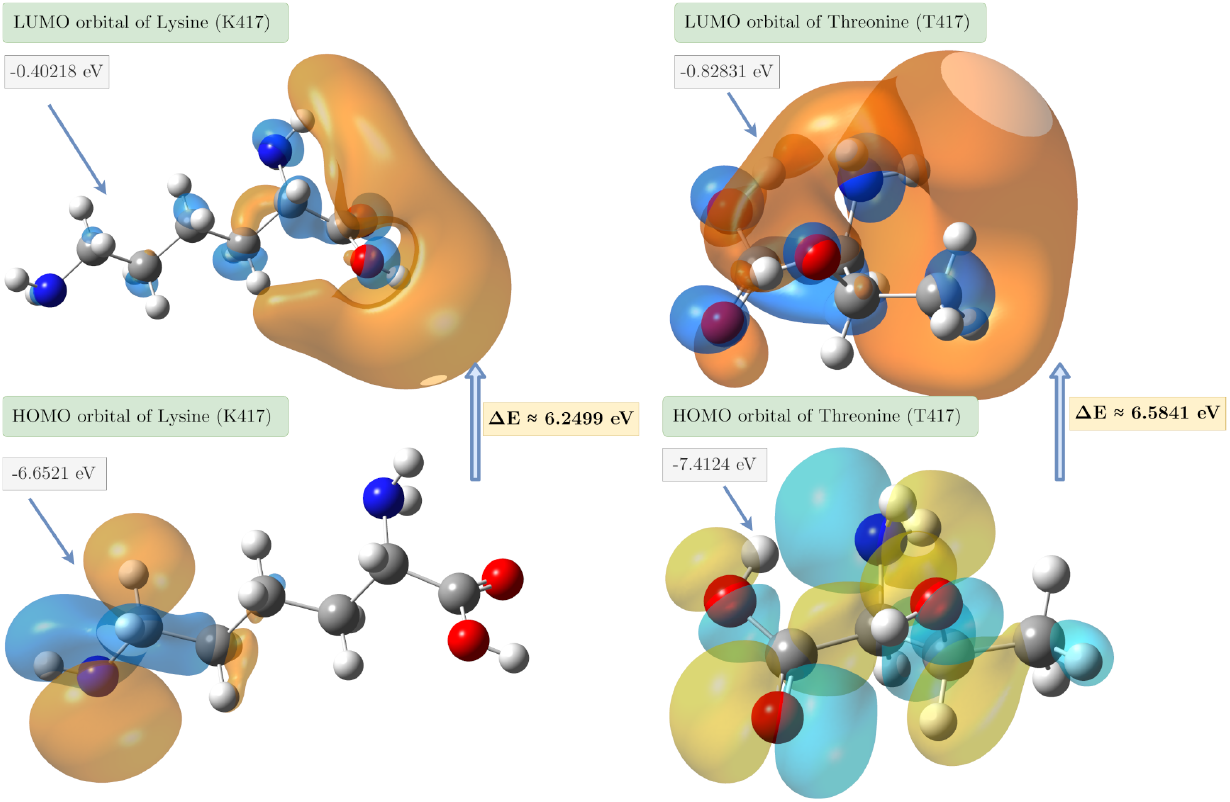
Comparison between the border orbitals of the amino acids Lysine and Threonine that constitute the K417T mutation of SARS-CoV-2.

**Figure 8.**
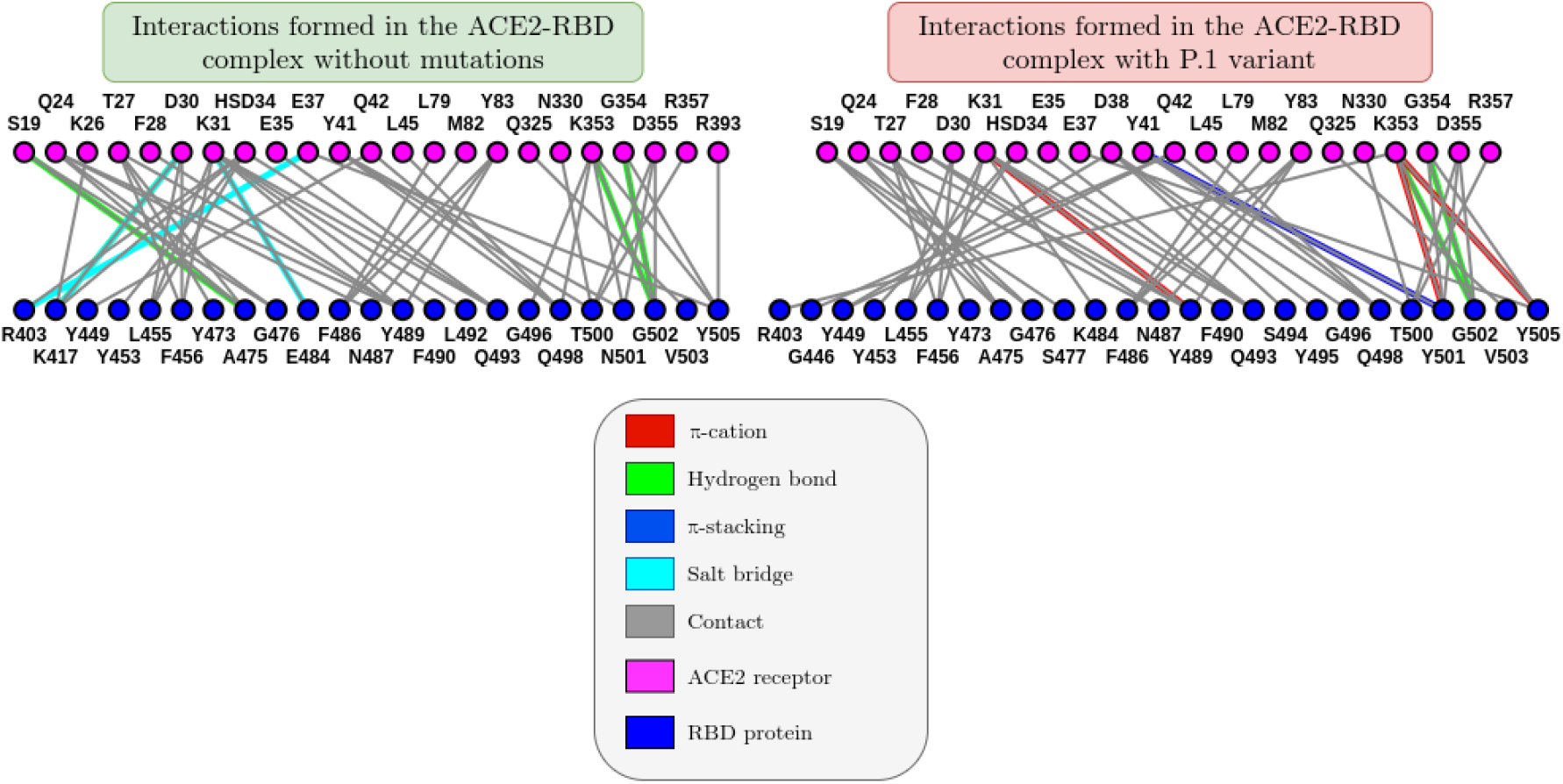
Comparison between the graphs of the interaction between the amino acids of the ACE2 receptor with those of the RBD protein referring to the last of the crystallographic structure PDB ID: 6M0J. The images were generated on the ICn3D online platform (https://www.ncbi.nlm.nih.gov/Structure/icn3d/).

We performed the structural alignment (see Figure 9) in relation to the *Cα* of the last frame obtained from the molecular dynamics referring to the P.1 line and the respective initial structure to understand the impacts of the temporal evolution. Thus, using the Schrödinger Maestro 2020-4 software, the alignment resulted in an atomic displacement with RMSD of 2.1203Å which therefore reflects relative structural changes at the global level as a result of This result was also corroborated by the PDBeFold Krissinel and Henrick (2004) platform with 1.320Å RMSD and by TM-Align with 2.05Å. For the simulation absent from mutations in relation to the initial frame, where there was an RMSD of 1.8135Å, when using TM-Align the result was 1.81Å e through PDBeFold it was 1.256Å. These results are therefore an indication that P.1 presented more expressive conformational fluctuations in the ACE2-RBD interaction when compared to the structure without any mutations, although P.1 is still characterized by greater stability.

**Figure 9.**
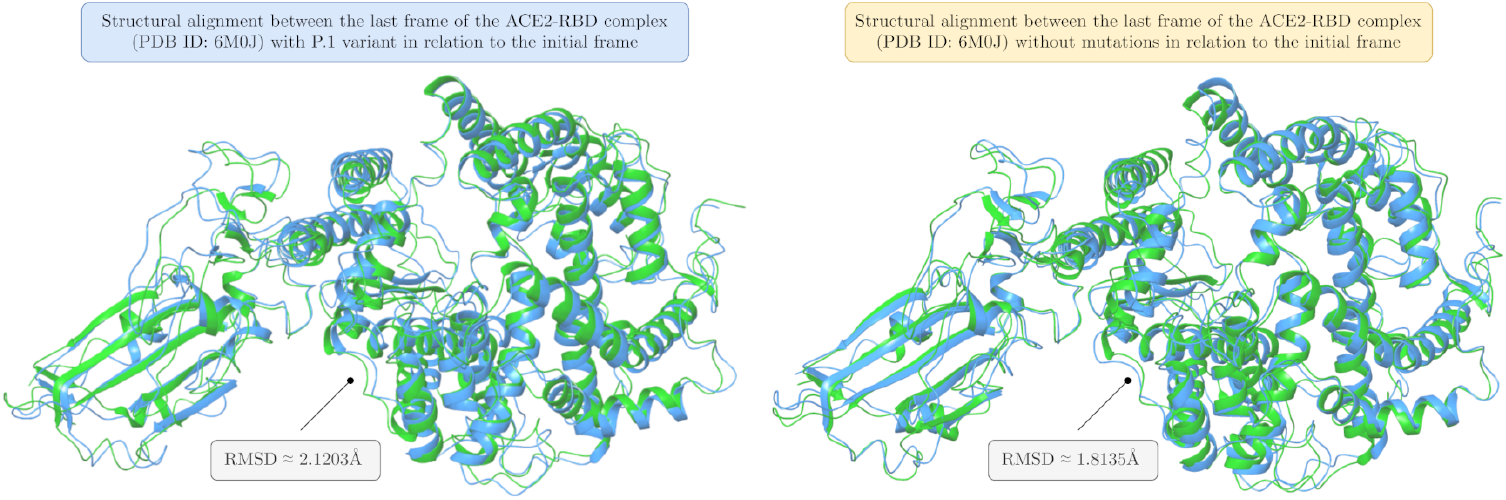
Comparison between the structural alignments for the last frame of the P.1 variant and the wild-type structure, with the respective initial frame for the ACE2-RBD complex (PDB ID: 6M0J). The image was made using the Schrödinger Maestro 2020-4 software.

Through RMSF analyzes it is possible to know the fluctuation variation around a specific residue (see Figure 10), thus allowing to understand the theoretical impacts of the mutations that make up the B.1.1.28/P.1 lineage. The K417T mutation indicated RMSF fluctuations in the Spike protein around 0.857Å, being higher than that affected the P.2 variant where K417N was 0.725Å. While N501Y in the amount of 0.864Å and finally E484K with 1.26Å. In terms of fluctuations, the E484K residue that emerged in South Africa was the most worrisome because it induced greater conformational changes in the Spike protein, although may be only because it is a naturally more unstable position and perhaps that is why it is more susceptible to mutations. In addition, higher RMSF values were found in the loops, possibly because they are not very defined regions. The greater stabilization in the ACE2-RBD complex as a result of certain mutations may be an explanation of why the virus has been following a convergent evolution as already reported in some experimental studies (Bobay, O’Donnell, & Ochman, 2020; Hodcroft et al., 2021; Kemp et al., 2021; Lam et al., 2020), although the causes until still unknown. Through the simulations of this work, we conclude that there is a tendency towards greater structural stability in the most frequent mutations. In other words, the mutations that have been recurring in several strains such as E484K tend to have greater stability.

**Figure 10.**
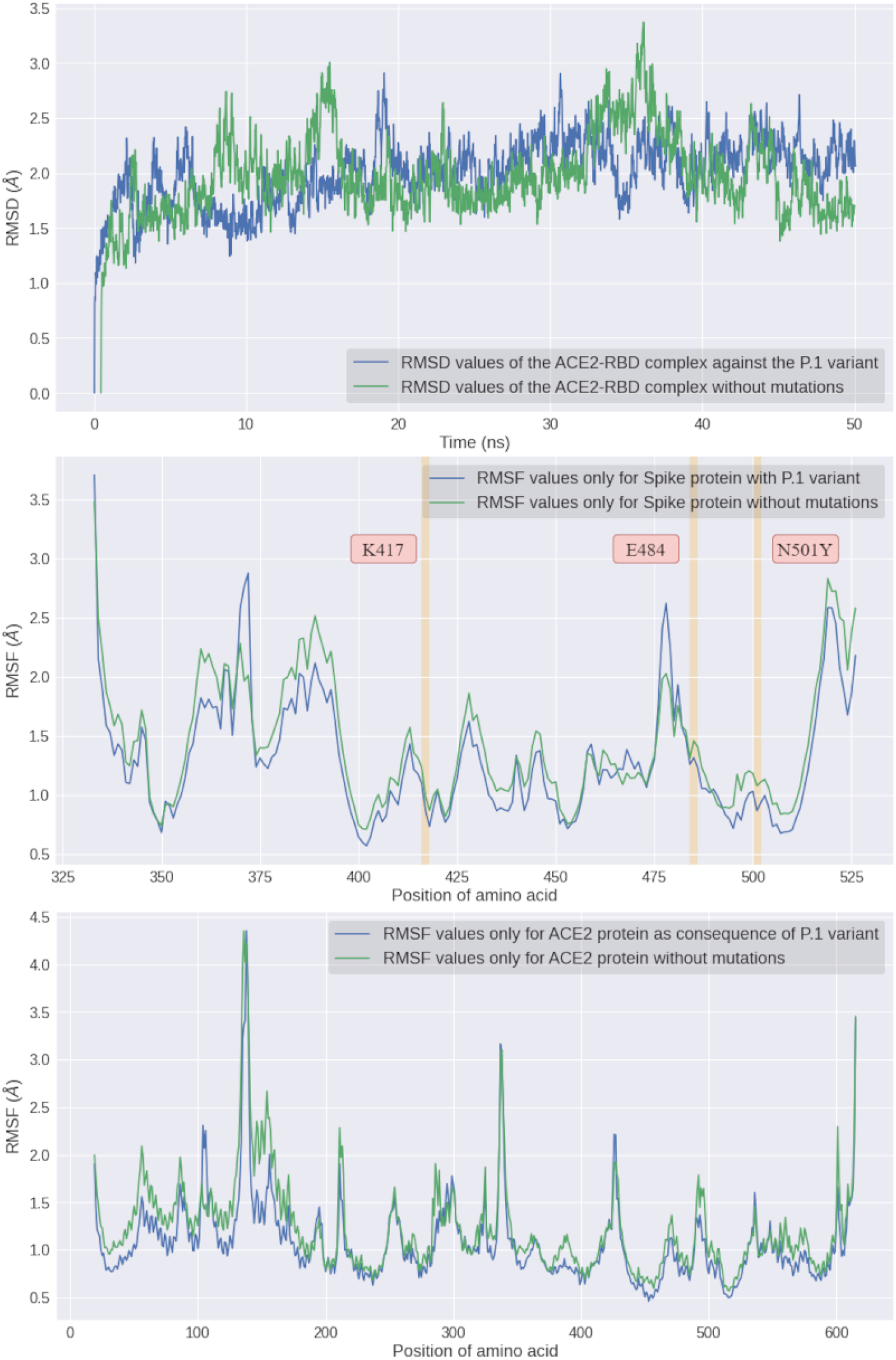
Comparative analysis between RMSD and RMSF fluctuations in relation to *Cα* with and without the Brazilian lineage B.1.1.28/P.1.

With the help of the RMSD diffusion map (see Figure 11), we can see that in the face of variant P.1 the map was relatively darker, and therefore more stable due to lower RMSD values, which favors ACE2-RBD interaction as a result of mutations.

**Figure 11.**
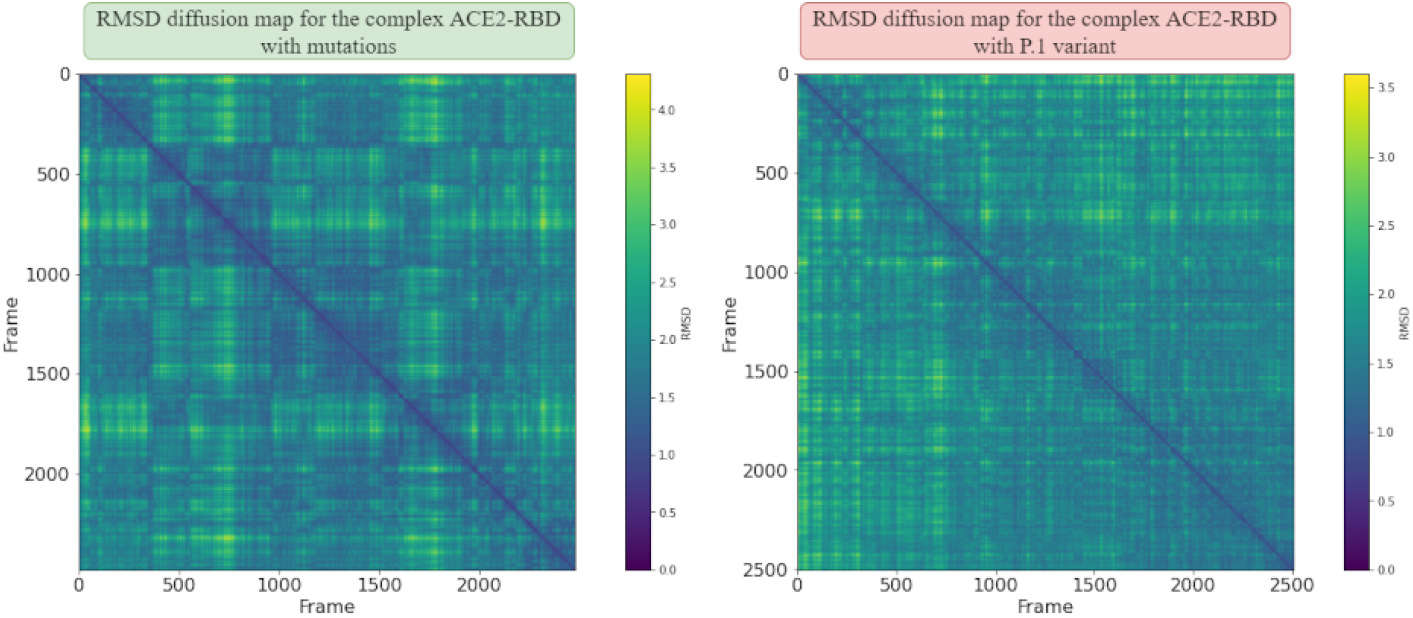
Scatter map of RMSD values over 50*ns* for the ACE2-RBD complex (PDB ID: 6M0J). A qualitative comparison of conformational changes in the face of the P.1 variant and in the structure absent mutations is presented.

When analyzing the ACE2-RBD complex without any mutations, we realized that the RMSF fluctuations (see Figure 3.2.1) for residue K417 was approximately 0.976Å, compared to residue N501 it was 1.08Å and lastly the amino acid E484 was 1.31Å. In the absence of mutations, the RMSF peak was around 2.83Å, but when the P.1 strain was analyzed there was an increase in the maximum fluctuation to 2.88Å. In general, although the fluctuation differences may be partly a consequence of a new velocity distribution associated with each simulation performed, the mutations indeed contributed to an increase in conformational changes in some very specific regions. This is because, when analyzing only the average of fluctuations (see Table 5), a lower RMSD is noticeable in P.1 and therefore greater stability. Consequently, our hypothesis is that the increase in transmissibility it may be correlated with greater affinity in the interaction of the ACE2-RBD complex as a consequence of the highest stability. It seems paradoxical that although P.1 presents comparatively larger peaks of fluctuations, the average RMSD was lower compared to the structure absent from mutations. Despite everything, it is consolidated in the literature that in the protein-protein interaction, greater stabilization is the main factor for increasing affinity. Therefore, the possible explanation for the highest peak is an indication that some very specific regions tend to suffer more from the impact of mutations, although at the global level there are fewer fluctuations. Finally, at the same time that the Spike protein becomes more flexible in some regions as a result of P.1, it may be indicative of greater exposure to neutralizing antibodies, although there may be a decrease in the affinity of interaction due to the RMSF instability of the system.

**Table 3.**
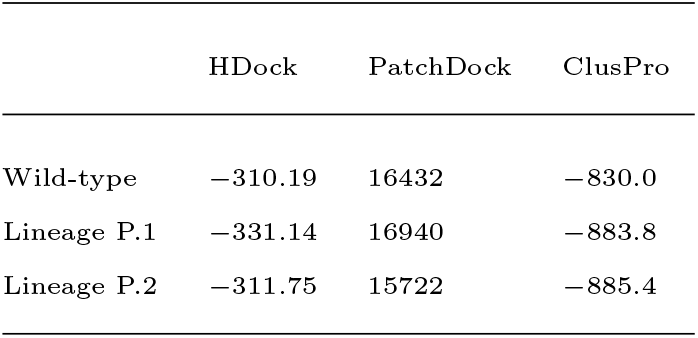
Results of protein-protein docking for the ACE2-RBD complex (PDB ID: 6M0J) from the last frame of the MD simulations, and whose affinity was estimated with the HDock, PatchDock and ClusPro tools.

|  | HDock | PatchDock | ClusPro |
| --- | --- | --- | --- |
| Wild-type | -310.19 | 16432 | -830.0 |
| Lineage P.1 | -331.14 | 16940 | -883.8 |
| Lineage P.2 | -311.75 | 15722 | -885.4 |

**Table 4.** Results of electronic descriptors obtained by structural optimization at the level of DFT theory with the Gaussian base 6 − 311*G* + +(2*d*, 2*p*) and B3LYP hybrid function. All minimizations were made using the Gaussian 09W software.

| Term | Lysine | Threonine |
| --- | --- | --- |
| Minimum energy | -13529.8117eV | -11930.5981 |
| Dipole moment | 0.902509Debye | 4.266042Debye |
| $E_{HOMO}$ | -6.6521eV | -7.4124eV |
| $E_{LUMO}$ | -0.40218eV | -0.82831eV |
| $E_{HOMO-1}$ | -7.0959eV | -8.1397eV |
| $E_{LUMO+1}$ | -0.33633eV | -0.17959eV |
| $\Delta E$ | 6.2499eV | 6.5841eV |
| Electron affinity | 0.40218eV | 0.82831eV |
| Ionization affinity | 6.6521eV | 7.4124eV |
| Chemical hardness ($\eta$) | 3.1250eV | 3.2920eV |
| Chemical softness ($\eta^{-1}$) | 0.16000eV | 0.15188eV<sup>-1</sup> |
| Chemical potential ($\mu$) | -3.5271eV | -4.1203eV |
| Electrophilicity index ($\omega$) | 1.9905eV | 2.5785eV |
| Electronegativity ($\chi$) | 3.5271eV | 4.1203eV |
| Nucleophilicity index ($N$) | 2.7165eV | 1.9562eV |

**Table 5.** Comparison between the P.1 and P.2 variants in relation to the average values of some parameters resulting from the molecular dynamics in the range of 50*ns* for ACE2-RBD (PDB ID: 6M0J) that quantify structural changes in the Spike RBD region in the interaction with ACE2.

| Analysis | Without mutations | Lineage P.1 | Lineage P.2 |
| --- | --- | --- | --- |
| Average RMSD | $(1.97 \pm 0.36)\text{\AA}$ | $(2.01 \pm 0.29)\text{\AA}$ | $(1.81 \pm 0.34)\text{\AA}$ |
| Average RMSF | $(1.28 \pm 0.51)\text{\AA}$ | $(1.13 \pm 0.46)\text{\AA}$ | $(1.33 \pm 0.76)\text{\AA}$ |
| $H \cdots$ bonds | $197 \pm 11$ | $201 \pm 11$ | $203 \pm 11$ |
| Average SASA | $(380.41 \pm 3.47)\text{nm}^2$ | $(377.64 \pm 3.66)\text{nm}^2$ | $(373.35 \pm 3.32)\text{nm}^2$ |
| Native contacts | $0.9903 \pm 0.0017$ | $0.9892 \pm 0.0016$ | $0.9877 \pm 0.0019$ |
| Average Rg | $(31.557 \pm 0.296)\text{\AA}$ | $(31.573 \pm 0.183)\text{\AA}$ | $(31.48 \pm 0.18)\text{\AA}$ |

Through analyzes of the average RMSD value (see Table 5), the absence of mutations in the ACE2-RBD complex induced approximately (1.97 ± 0.36)Å in atomic displacement and average RMSF fluctuations of (1.28 ± 0.51)Å. On the other hand, in the presence of P.1 lineage mutations, the average RMSD decreased to (2.01 ± 0.29)Å, while the average RMSF value decreased to (1.13 ± 0.46)Å. Thus, the higher RMSD in the absence of mutations can be associated with a greater probability of detachment of the Spike protein with the ACE2 receptor interface. In addition, *α*-helices for being present in naturally unstable positions were the ones that presented the highest conformational fluctuations, and that also suffered the direct effects of the mutations. After identification by the VMD software, we noticed that the Asn370 residue in the RBD protein showed an RMSF fluctuation of 2.58Å in the structure containing the P.1 lineage, but in the absence there was an increase in stability to 2.28Å. It is noteworthy that this region was the one that showed the greatest fluctuation among all amino acids regardless of the presence of mutations. Therefore, these data possibly reflect an increase in the ACE2-RBD interaction due to lower RMSD displacements as a result of the multiple mutations that make up the P.1 strain.

Regarding the analysis of the formation of Hydrogen bonds over the ACE2-RBD simulation (see Figure 13), it was found that the P.1 strain had an average (see Table 5) of 201 ± 12 hydrogen bonds and in the absence there was a decrease to 197 ± 11 on average bonds. The number of hydrogen bonds at the ACE2-RBD interface was shown to be lower in P.1 compared to the structure absent from mutations, since in P.1 there were 31 interactions at the interface while in the wild-type structure 34 interactions were formed throughout the simulation. However, this does not allow us to conclude that the structural stability of P.1 is less. This is because only 2 (two) hydrogen interactions are predominant in the complex, where for P.1 we have the formation of ACE2: Gly502 → RBD: Lys353 with an incidence of 58.04% in addition to ACE2: Tyr83 → RBD: Asn487 with 51.07%. When compared to the missing structure of mutations and frequency it is lower where ACE2: TYR83 → Asn487 with 54.26% and ACE2: Lys417 → RBD: Asp30 with 52.28%. Therefore, when only more frequently hydrogen bonds are analyzed, P.1 is predominant in its formation, which is an important factor for structural stability.

**Figure 12.**
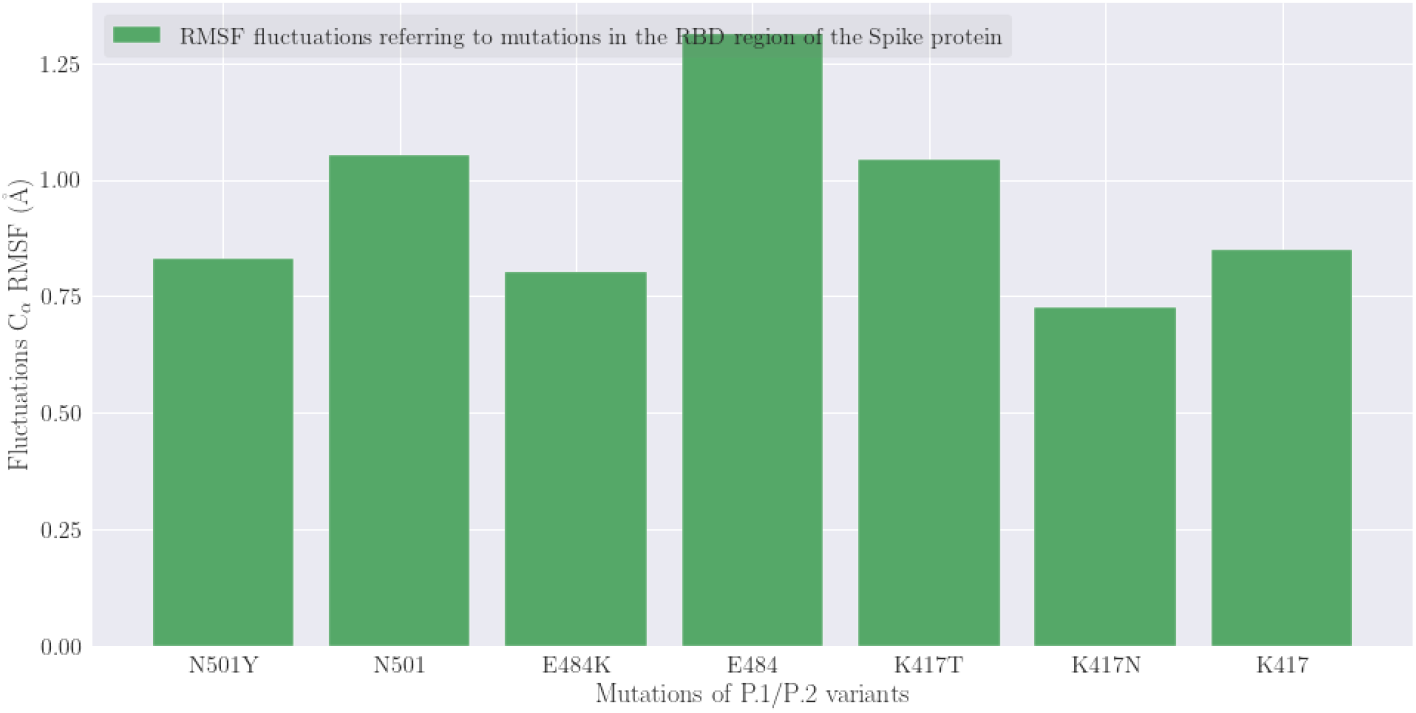
Individual comparisons of RMSF atomic fluctuations obtained from molecular dynamics in NAMD3 algorithm. Amino acids that were affected by the P.1/P.2 variants are presented as well as simulation results where no mutations were applied.

**Figure 13.**
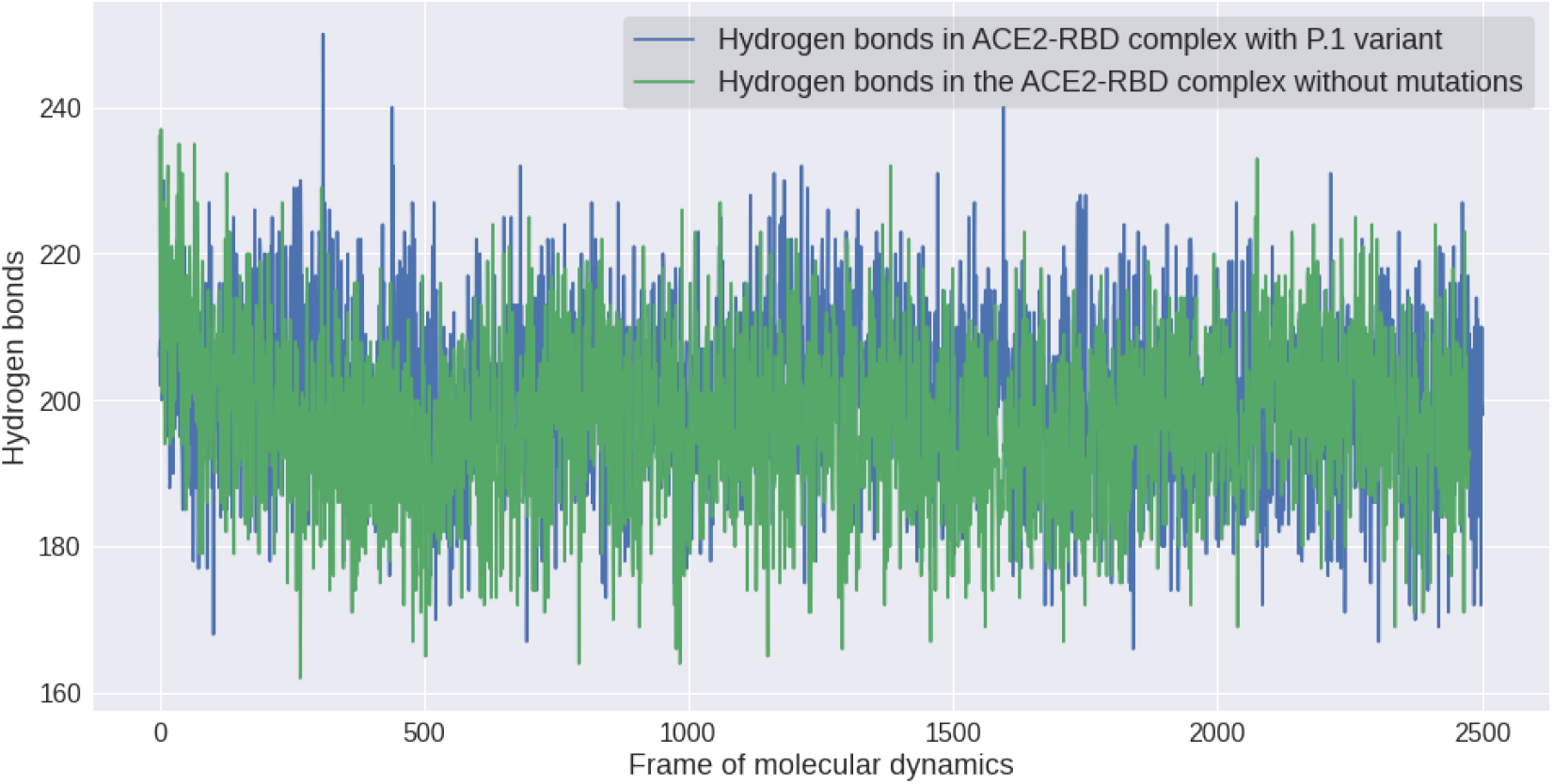
Impact on the formation of hydrogen bonds in the ACE2-RBD complex as a result of the P.1 lineage.

When analyzing the formation of saline bridges, it was noticed that only the wild-type structure formed 2 (two) interactions in the ACE2-RBD interface between the residues: RBD: Glu484 → ACE2: Lys31 and RBD: Lys417 → ACE2: Asp30. On the other hand, the structure containing P.1 has lost these interactions although formed only the interaction RBD:Lys484 → ACE2:Glu75. Consequently, we can conclude that Hydrogen bonds play a more fundamental role in the stabilization of P.1 to the detriment of saline bridges. Therefore, the formation of more chemical interactions, especially hydrogen bonds, therefore seems to increase the structural stability of P.1. In summary, there seems to be a favoring of this lineage so that there is a greater ACE2-RBD interaction.

A larger area accessible to the solvent reduces the chances of interactions between the ACE2 receptor and the virus, since Spike central amino acids would be clogged by the solvent. On the other hand, a lower SASA also represents an increase in hydrophobic interactions. In summary, the increase in the SASA value indicates protein expansion, and thus, the higher its value, the more unstable system becomes and the smaller the interaction becomes. When analyzing the total SASA value over time (see Figure 14) the differences were statistically significant where P.1 presented an average at (377.64 ± 3.66)*nm*^2^ differing only (−2.77*nm*^2^) compared to the structure absent from mutations (see Table 5). This is therefore an indication that exposure to the solvent although was affected by the mutations that make up the P.1 strain, but that it does not play a central role in increasing ACE2-RBD affinity. From the joint analysis of RMSD, RMSF, SASA and Hydrogen bonds, it can be seen that mutations of the P.1 strain have stabilized the ACE2-RBD structure, although in some central residues a decrease in structural flexibility has been noted according to RMSF analysis.

**Figure 14.**
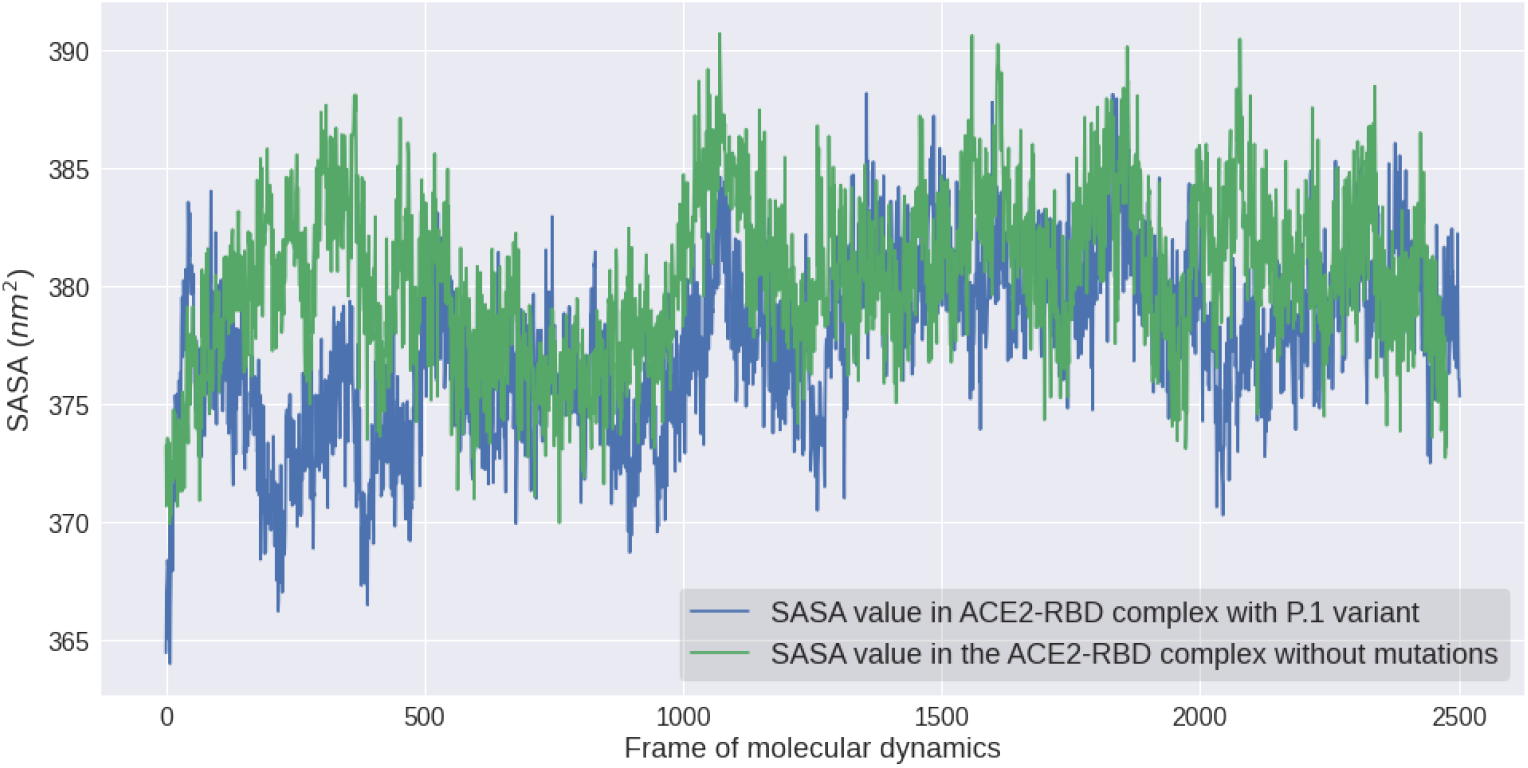
Comparison of the SASA values for exposure to the solvent of the ACE2-RBD complex (PDB ID: 6M0J) in view of the mutations that make up the P.1 strain.

Finally, it is important to highlight that 3 (three) analyzes corroborate the hypothesis of greater ACE2-RBD stability as a result of P.1 (see **??**), among them: Low mean RMSF values, greater formation of bonds of Hydrogen and lower solvent exposure measured by the SASA value. While only 2 (two) analyzes reflect the hypothesis of greater instability, namely: A smaller fraction of native contacts and smaller compaction measured by the Radius of Rotation. However, we must remember that the mean RMSF value is naturally more unstable in the face of more expressive peak values as a result of mutations, although in general we can indeed conclude that in fact stability is the main conclusion for changes in P.1 transmissibility and virulence.

A greater deviation from native contacts as a result of the P.1 lineage was noticeable in the ACE2-RBD complex (see Figure 15), decreasing from 0.9903±0.0017 to 0.9892± 0.0016. This therefore indicates notable and statistically significant structural changes (*p <* 0.05). Thus, these modifications may have a correlation to the appearance of a phenotypic characteristic where there is greater viral transmissibility in similar strains such as B.1.1.351 as reported in an important study (Faria et al., 2021).

**Figure 15.**
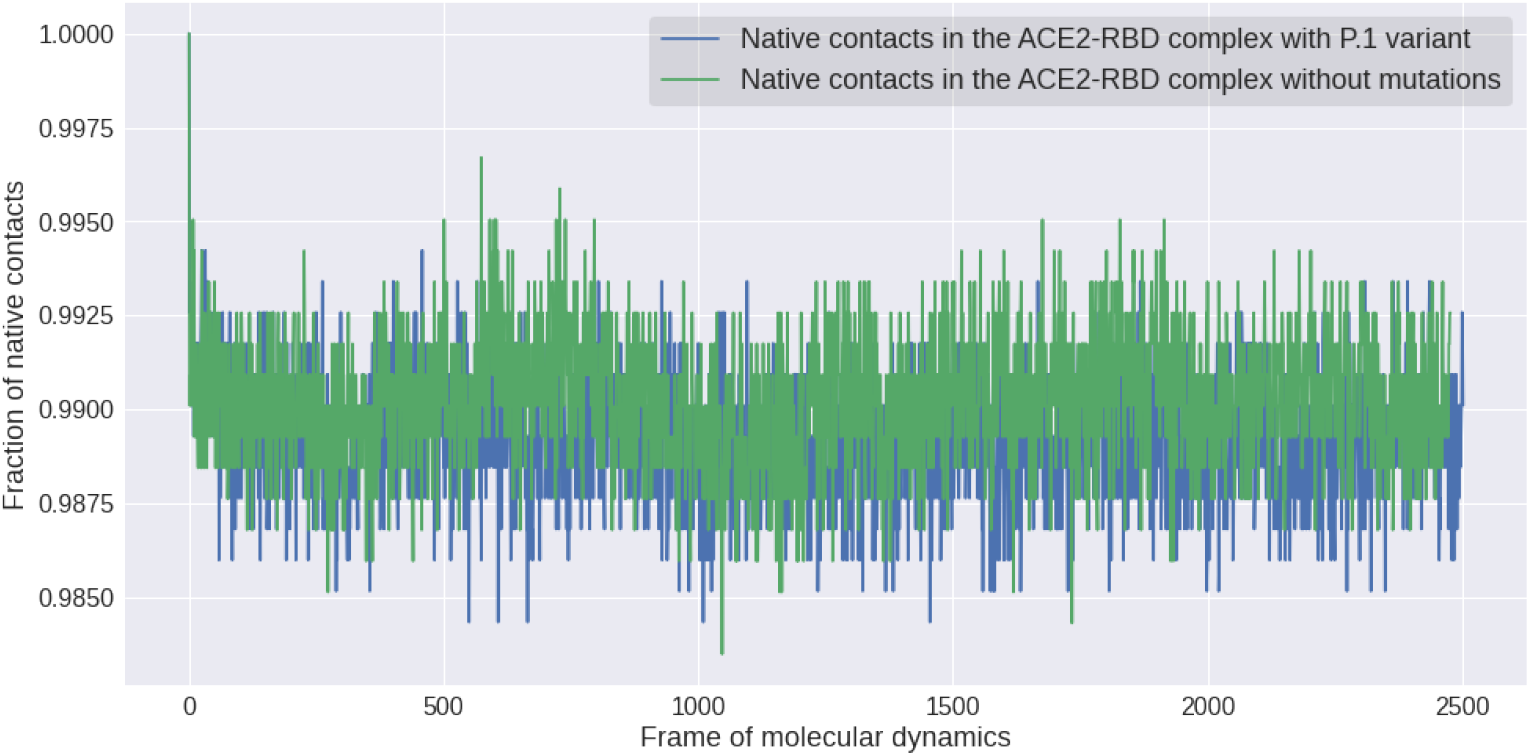
Comparison between the fractions of native *Q*(*t*) contacts in the ACE2-RBD complex (PDB ID: 6M0J) as a result of the mutations that constitute the P.1 lineage.

In terms of compaction of the ACE2-RBD complex (see Figure 16) we can see that the P.1 lineage induced greater packaging along the interaction described by the average Radius of Gyration (Rg) of (31.573 ± 0.183)Å when analyzing all Cartesian axes. On the other hand, in the absence of mutations, there was an increase in Rg to (31.557 ± 0.296). When performing a statistical comparison using the Student’s t-test, the Pearson’s coefficient value was *p <* 0.05 within the 95% confidence interval, thus showing a statistically significant difference between the groups analyzed. Although the statistical approach was important to minimize subjectivity in the interpretation of the results, it is important to note that the *p* coefficient cannot be seen as something absolute or that the differences are necessarily statistically significant (Wasserstein, Schirm, & Lazar, 2019). In spite of everything, the current paradigm is that strains similar to P.1 really generated an increase in transmissibility and lethality (Faria et al., 2021), and consequently we seek to see this in the results through the greater ACE2-RBD affinity, compaction and system stabilization as a consequence of P.1 lineage.

**Figure 16.**
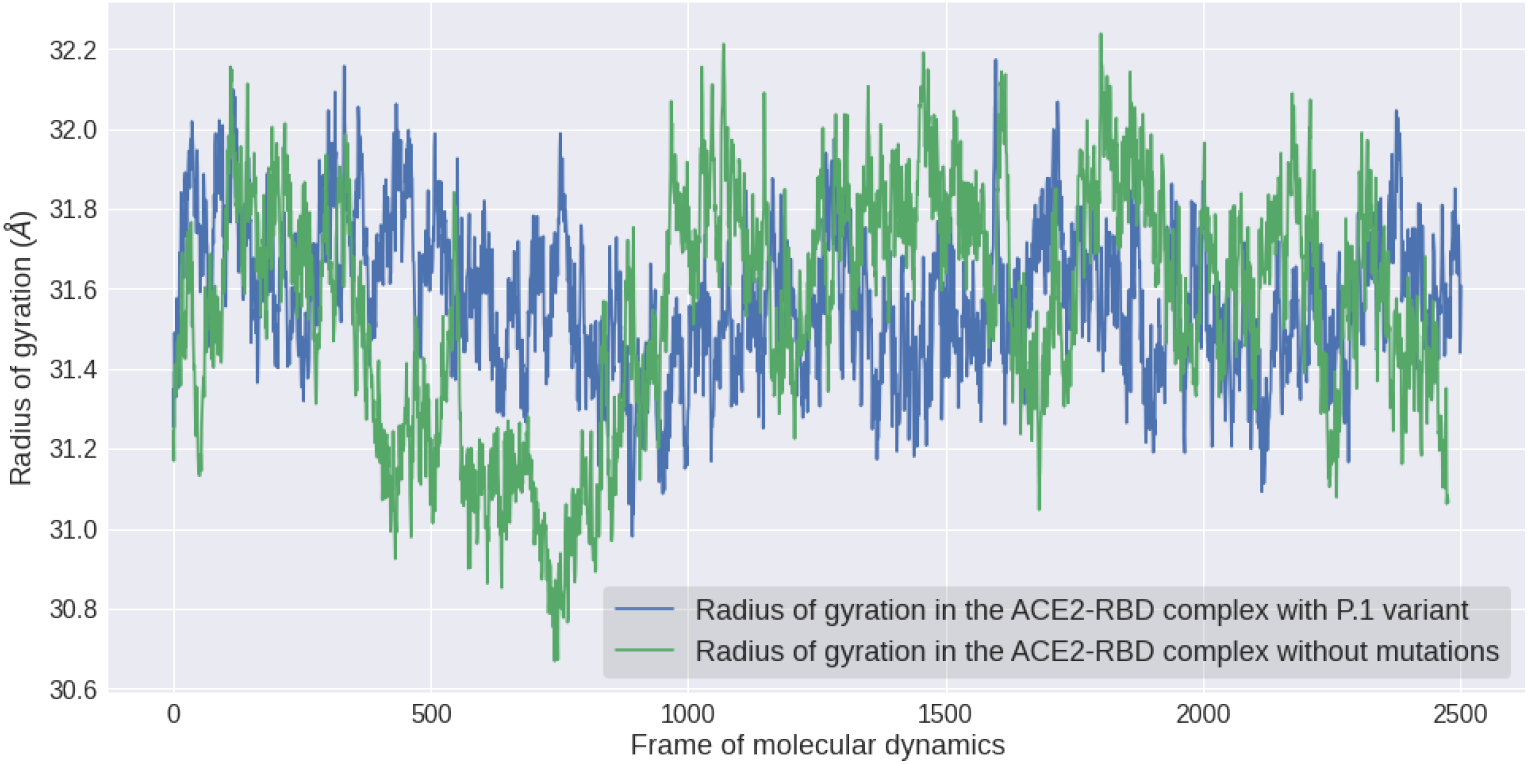
Comparative plot between the radius of gyration (Rg) values in the presence and absence of P.1 lineage. We noticed that the simulations in GROMACS showed greater compaction and therefore less turning radius compared to the P.1 variant.

Through the simulations of molecular dynamics in the NAMD 3 algorithm, it was also possible to understand the impacts of the P.1 strain on the effectiveness of neutralizing antibody in the interaction with the Spike protein (RBD). The antibody studied, P2B-2F6, is naturally produced by the human organism in face of SARS-CoV-2 infection. Therefore, it is not obtained synthetically by the biotechnology industry, since the costs of manufacturing monoclonal antibodies are extremely high. From the individual analysis of RMSF fluctuations associated with each residue, the impact of the P.1 strain was realized so that the E484K mutation in the RBD + P2B-2F6 antibody complex (PDB ID: 7BWJ) generated an increase in instability of (1.59 ± 0.71)Å → (1.43 ± 0.52)Å. As for the K417N mutation, there was the same behavior with the transition from 1.139Å → 1.543Å. Again these results were repeated for N501Y where the change was 1.653Å → 2.154Å. In this way, it seems that a characteristic of P.1 is precisely to induce less stability in the interaction with neutralizing antibodies, which reflects in less affinity of the interaction. Furthermore, it was noticed a decrease of 1 (one) saline bridge during the MD simulations, where the RBD + P2B-2F6 antibody complex (PDB ID: 7BWJ) lost the interactions between the residues RBD: Glu484 → Heavy Chain: Arg112 and RBD: Glu484 → Light Chain: Lys55 although it formed RBD: Lys484 → Light Chain: Glu52. This therefore reflects a possible decrease in the interaction of neutralizing antibodies due to greater instability in the antibody-antigen complex. Lastly, the alignment of the crystallographic structure of the RBD + P2B-2F6 antibody complex (PDB ID: 7BWJ) with the last frame resulting from the molecular dynamics with P.1, resulted in an RMSD value of 2.5636Å, which indicates substantial conformational changes. All graphical comparative analyses for RMSD, RMSF, Hydrogen bonds and others, for the RBD + P2B-2F6 complex (PDB ID: 7BWJ), are found as *Supplementary Material*.

In general, almost all comparisons were statistically significant, with the exception of the SASA value. In order to justify the validity of the t-test approach employed, the Shapiro-Wilk normality test was performed for the number of hydrogen bonds, as a test example, with a minimum confidence interval of 87%. Levene’s test was applied to verify equality between variances where *p >* 0.05, and therefore the inequality had to be considered throughout the hypothesis test. Consequently, we believe that the T-Test formalism proposal for the analysis of MD simulations is justified. Regarding the RMSD and RMSF data set, the confidence interval of 95% was adopted. Finally, we must note that the results of any computational simulation must always be seen with the perspective that the improvement of the force fields and algorithms is continuous, and therefore these results are provisional until corroborated by more in-depth theoretical studies or using experimental techniques. The differences between P.1 and will become clearer as we repeat the simulations or even in longer intervals for more conclusive results and greater reproducibility. Anyway, as in the Spike RBD region the only difference between the variants is the change from the K417T to K417N mutation, so it is expected that the structural changes are not as noticeable between the two variants. Therefore, similar conclusions are expected for the two strains that emerged in Brazil. Although molecular dynamics is a rigorous approach to study the interaction of biomolecules, it is necessary to calculate as many trajectories as possible to minimize statistical errors.

The most recent variant of concern appeared in India, being called B.1.617.2 (Delta), presenting mutations until then in regions not affected by old lineages, where we have in the RBD region: L452R, T478K. However, the same region that affected P.1 (E484K mutation), led to the emergence of another mutation, E484Q. This therefore reinforces the convergence of certain mutations, but raises questions because new amino acid positions have been affected.

### 3.3. Molecular dynamics with P.1 variant crystallography

Only very recently has a crystallography containing the P.1 variant been made available. Thus, in addition to MD simulations for computationally inserted mutations, we also found it necessary to simulate with experimentally characterized mutations. When we simulated from the P.1 variant itself crystallography (see Table 8), we can obtain conclusions more consistent with reality. Thus, it was noted that P.1 provided, in general, lower structural stability in the antibody-antigen interaction, which therefore reduces the interaction affinity. In summary, 3 (three) analyzes support the hypothesis of lower stability: Higher values of RMSD and RMSF fluctuations, in addition to the lower number in the fraction of native contacts.

**Table 6.** Comparison between the average values of some parameters of molecular dynamics referring to the simulation of the RBD region of the Spike protein with the neutralizing antibody P2B-2F6 (PDB ID: 7BWJ). All these results refer to simulations in the range of 30*ns* performed on the NAMD3 algorithm.

| Analysis | Without mutations | Lineage P.1 | Lineage P.2 |
| --- | --- | --- | --- |
| Average RMSD | $(3.75 \pm 0.85)\text{\AA}$ | $(2.48 \pm 0.37)\text{\AA}$ | $(2.40 \pm 0.56)\text{\AA}$ |
| Average RMSF | $(1.59 \pm 0.71)\text{\AA}$ | $(1.43 \pm 0.52)\text{\AA}$ | $(1.75 \pm 0.56)\text{\AA}$ |
| Average $H \cdots$ | $160 \pm 10$ | $158 \pm 9$ | $155 \pm 9$ |
| Average SASA | $(309.49 \pm 314)\text{nm}^2$ | $(306.47 \pm 2.50)\text{nm}^2$ | $(313.55 \pm 380)\text{nm}^2$ |
| Native contacts | $0.981 \pm 0.003$ | $0.978 \pm 0.003$ | $0.980 \pm 0.003$ |
| Average Rg | $(33.60 \pm 0.42)\text{\AA}$ | $(32.95 \pm 0.26)\text{\AA}$ | $(33.51 \pm 0.30)\text{\AA}$ |

**Table 7.** Comparison of statistical significance between the P.1 variant and the wild-type structure from NAMD3 simulations. Regarding the RMSD and RMSF data set, the 95% confidence interval was adopted.

| Complex | PDB ID | p-value |  |  |  |  |  |
| --- | --- | --- | --- | --- | --- | --- | --- |
| | | *RMSD* | *RMSF* | Hydrogen Bonds | SASA | $Q(t)$ | $R_g$ |
| ACE2-RBD | 6M0J | $p < 0.05$ | $p < 0.05$ | $p < 0.05$ | $p < 0.05$ | $p < 0.05$ | $p < 0.05$ |
| P2B-2F6 | 7BWJ | $p < 0.05$ | $p < 0.05$ | $p < 0.05$ | $p < 0.05$ | $p < 0.05$ | $p < 0.05$ |

**Table 8.** Analysis of molecular dynamics simulations of the antibody-antigen complex with P.1 (PDB ID: 7NXB) and without (PDB ID: 7NX6) using GROMACS software.

| Analysis | Without mutations | Lineage P.1 |
| --- | --- | --- |
| Average RMSD | $2.07 \pm 0.33$ | $2.90 \pm 0.62$ |
| Average RMSF | $1.21 \pm 0.52$ | $1.74 \pm 0.80$ |
| $H \cdots$ bonds | $158 \pm 10$ | $161 \pm 10$ |
| Average SASA | $(301.76 \pm 2.39)nm^2$ | $(299.85 \pm 3.20)nm^2$ |
| Native contacts | $0.9794 \pm 0.0030$ | $0.9785 \pm 0.0032$ |
| Average Rg | $31.28 \pm 0.19$ | $30.95 \pm 0.23$ |

#### 3.3.1. MM-PBSA decomposition results

The Van der Waals interaction and electrostatic energy were estimated in addition to the solvation energy. Thus, will help us to better understand the theoretical causes of the greater stability of the ACE2-RBD complex as a result of P.1 mutations. Throughout the discussion of the results (see Table 10), we focused on the terms that make up the ⟨*E*_*MM*_⟩ energy between the RBD chain and the ACE2 receptor, instead of the Δ*G*_*binding*_ because it would require accurate estimation of the entropic term. We consider that the internal energy component of molecular mechanics did not make a significant contribution, therefore approaching zero. The favorable formation and greater stability of the ACE2-RBD complex as a function of P.1 may be correlated mainly with even smaller terms of the energy of Van der Waals ⟨*E*_*V dW*_⟩. Apparently, Van der Waals contribution (see Figure 17) has been playing a central role in a greater formation of intermolecular bonds in the complex containing P.1. Finally, even with such diverse MD methodologies, whether by NAMD or GROMACS, the hypothesis that Van der Waals interactions predominate in the P.1 variant has been supported.

**Table 9.**
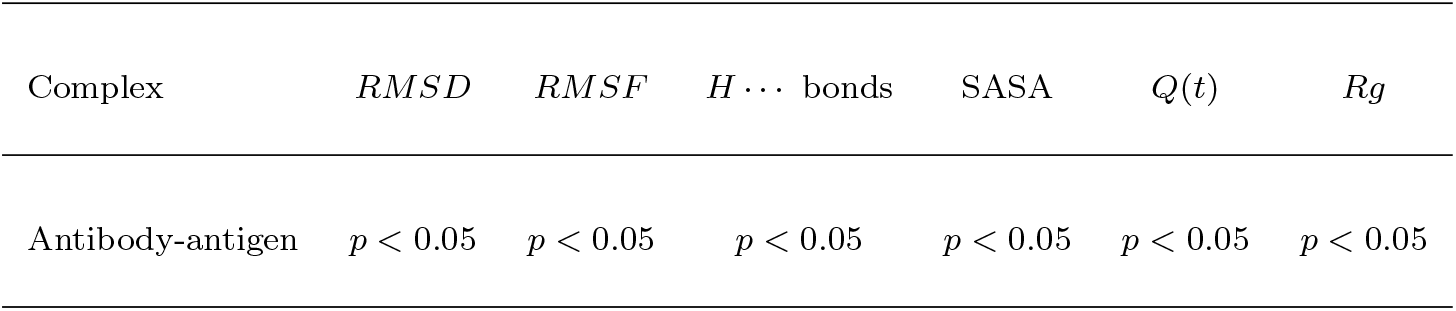
Comparison of statistical significance between the P.1 variant and the wild-type structure of GRO-MACS simulations. For the RMSD and RMSF dataset, the confidence interval 95% was adopted.

**Table 10.** All MM-PB(SA) energy decomposition values refer to the ACE2-RBD complex (PDB ID: 6M0J) in 50*ns* simulations with NAMD3. The energy values are in the unit *kcal · mol*^−1^. The term for *E*_*int*_ covalent interactions was considered to be approximately zero in the analyzes. In the free energy calculations of Δ*G* it was considered that the entropy variation Δ*S* was approximately zero and consequently Δ*G ≈* Δ*H*.

| Term | Without mutations | Lineage P.1 | Lineage P.2 |
| --- | --- | --- | --- |
| $\langle E_{elec} \rangle$ | $-178.32 \pm 40.88$ | $-62.69 \pm 32.53$ | $-45.91 \pm 25.62$ |
| $\langle E_{VDW} \rangle$ | $-67.98 \pm 5.30$ | $-70.19 \pm 4.97$ | $-74.15 \pm 4.91$ |
| $\langle E_{MM} \rangle$ | $-246.30 \pm 41.73$ | $-132.87 \pm 32.85$ | $-120.06 \pm 26.43$ |
| $G_{solvation}^{polar}$ | $-2753.57$ | $-2606.98$ | $-2570.16$ |
| $\langle G_{solvation}^{apolar} \rangle$ | $+207.10 \pm 1.88$ | $+205.60 \pm 1.98$ | $+203.28$ |
| $G_{solvation}^{total}$ | $2546.47$ | $2401.38$ | $-2366.88$ |
| Enthalpy | $2792.77$ | $2534.25$ | $-2486.94$ |
| $\Delta G$ | $-882.47$ | $-759.22$ | $-269.92$ |

**Figure 17.**
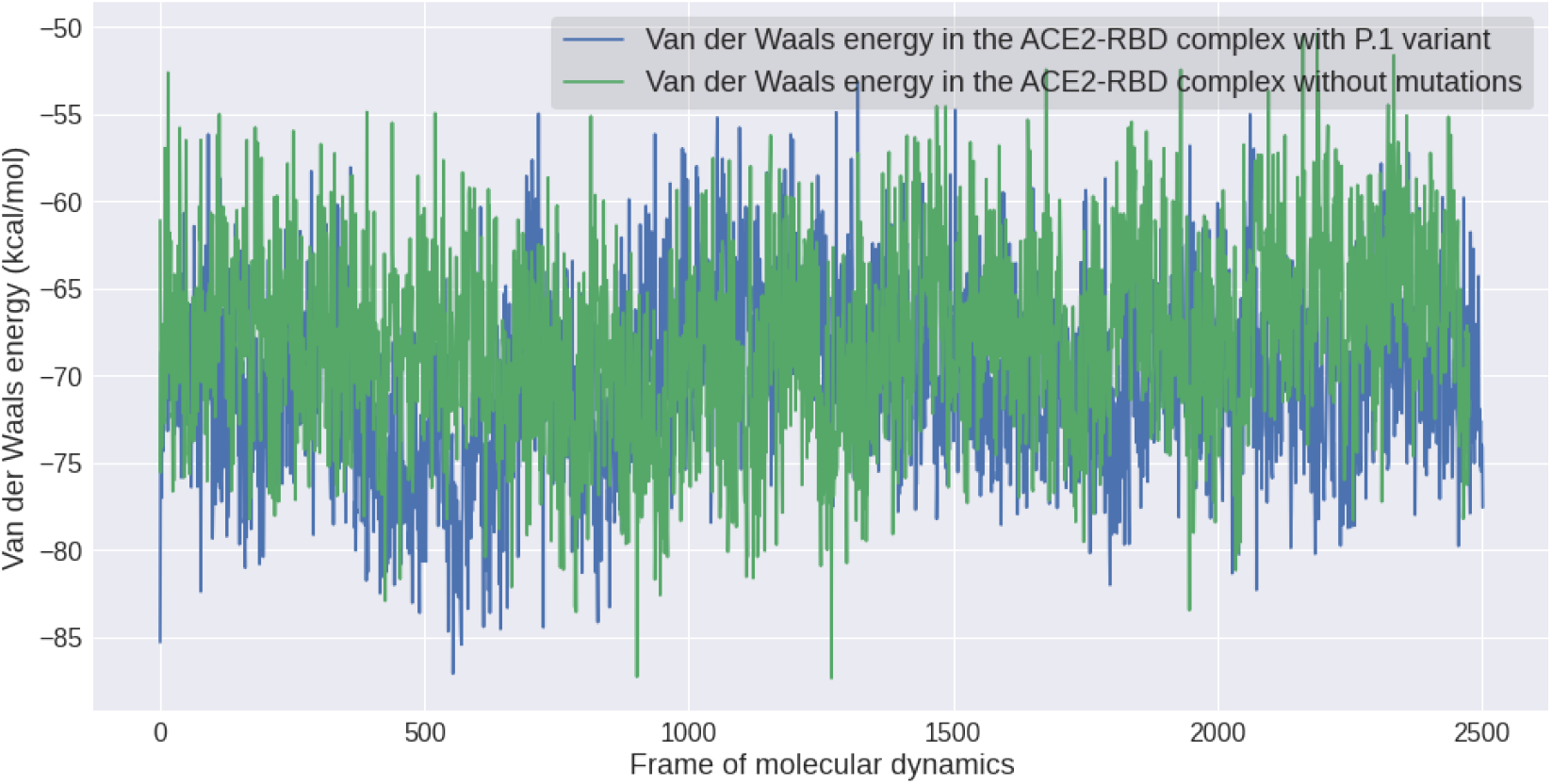
Study of the P.1 strain on the energy of Van der Waals ⟨*E*_*V dW*_⟩ on the interaction with the ACE2-RBD complex (PDB ID: 6M0J).

In terms of polar solvation, the impact of the P.1 strain was more notable compared to non-polar solvation where the difference was not significant. However, the polar solvation free energy *G*_*PB*_ favored the interaction of the wild-type complex much more, which could explain the greater instability and greater interaction with the surrounding environment instead of the ACE2 cell receptor. The unexpected results for electrostatic interactions may be a consequence of the numerous fluctuations in the energy ⟨*E*_*elec*_⟩ that characterized the wild-type structure ranging from −319.6277 *kcal · mol*^−1^ up to −80.3537 *kcal · mol*^−1^, and therefore increasing the instability of the ACE2-RBD system. Therefore, Van der Waals interactions are the most prevalent when comparing energy data between the complex containing and absent P.1 mutations. After estimating the molecular energy and solvation, it is finally possible to obtain the enthalpy *H* associated with the system. Thus, we realized that the P.1 strain unexpectedly had the less favorable enthalpic term compared to the reference structure, suggesting that greater ACE2-RBD interaction is correlated to less negative enthalpy but that it will supposedly reflect in a more favorable and therefore greater Δ*H* and spontaneity of Δ*G*_*binding*_. Finally, we must highlight that the energy values presented are very dependent on the methodology used. Consequently, numerical values should only be seen in a qualitative way for the impact of P.1 instead of absolute quantitative.

Firstly as a result of the ACE2-RBD interaction being a temperature-dependent process as discovered in the research (He, Tao, Yan, Huang, & Xiao, 2020) so the premises Δ*H <* 0 and Δ*S <* 0 are possibly true. Furthermore, it is known that Coronaviruses tend to spread with a higher incidence, even more negative values of Δ*G <* 0, due to the seasonality in winter. Consequently, we can conclude that the P.1 lineage, being more stable, will have less difficulty in overcoming the entropic penalty because greater affinity leads to a reduction in conformational entropy. This stems from the equation that describes the free binding energy between two proteins: Δ*G* = Δ*H* − *T ·* Δ*S*. As the term −*T ·* Δ*S* becomes even more positive as a result of less structural stability, the free energy of binding tends to be less spontaneous and the affinity of interaction becomes less expressive with the increase of temperature. In other words, the entropic penalty of free energy decreases in proportion to structural stability. The increased compaction of the ACE2-RBD complex as a result of the P.1 strain could also make the enthalpy variation more negative, which will lead to a greater propensity for proteolytic activation of the virus. Unexpectedly, the results for the ACE2-RBD complex, although they indicated greater structural stability, did not necessarily imply greater interaction affinity. This is because, the structure without mutations, strangely, presented a Δ*G ≈* −882.47*kcal · mol*^−1^ while against the P.1 variant it was less favorable with Δ*G ≈* −759.22*kcal · mol*^−1^. Although these results seem totally contradictory, we must remember that even consolidated prediction software such as Schrodinger Maestro, came to present Δ of affinity and stability with opposite signs to each other.

From the analysis of the vibrational modes of the last MD frames (see Table 12), we can see that the P.1 variant provided a smaller Δ*S* compared to the wild-type structure when analyzing the ACE2-RBD complex. This reflects in greater structural compaction, as the entropic contribution becomes smaller, and thus the interaction with P.1 becomes superior. It should be noted that as Δ*S* is greater, the interaction energy Δ*G* is impaired. Consequently, the hypothesis of greater structural stability has been confirmed by numerous molecular dynamics analyses, being the central feature of the P.1 variant.

**Table 11.** All MM-PB (SA) energy decomposition values refer to the P2B-2F6 neutralizing antibody in complex with RBD (PDB ID: 7BWJ) in 30*ns* simulations with NAMD3. The energy values are in the unit *kcal·mol*^−1^. Throughout the analyzes for the antibody-antigen complex, only the light chain (L) of the antibody was considered when interacting with the RBD. The term for *E*_*int*_ covalent interactions was considered to be approximately zero in the analyzes.

| Term | Without mutation | Lineage P.1 | Lineage P.2 |
| --- | --- | --- | --- |
| $\langle E_{elec} \rangle$ | $-10.67 \pm 17.23$ | $-59.51 \pm 27.41$ | $+4.43 \pm 17.75$ |
| $\langle E_{vdW} \rangle$ | $-8.37 \pm 2.53$ | $-5.66 \pm 3.92$ | $-3.82 \pm 4.07$ |
| $\langle E_{MM} \rangle$ | $-19.05 \pm 16.67$ | $-65.17 \pm 28.66$ | $+0.62 \pm 18.78$ |
| $G_{solv}^{polar}$ | -1319.65 | -1381.02 | -1367.02 |
| $\langle G_{solv}^{apolar} \rangle$ | +168.67 | $167.03 \pm 1.36$ | +170.87 |
| $G_{solv}^{total}$ | -1150.98 | -1213.99 | -1196.15 |
| *Enthalpy* | -1170.03 | -1279.16 | -1195.53 |

**Table 12.** Results of Δ*S*(*kcal · mol*^−1^ *· K*^−1^) for the last frame obtained by the algorithm ENCoM Frappier and Najmanovich (2014).

|  | ACE2-RBD | P2B-2F6 |
| --- | --- | --- |
| Without mutations | -19685.11 | 16048.60 |
| Lineage P.1 | -19681.55 | 16061.63 |
| Lineage P.2 | -19714.74 | 16032.13 |

According to the deformation energy analyzes present in the Figure 18, we noticed that in the face of the P.1 variant the mean value was 106522 ± 29008 while in the absence of mutations there was an increase to 107347 ± 28400. Therefore, this is an important indication that P.1 makes the Spike protein more stable, which is of concern as it increases the probability of interaction with the ACE2 receptor.

**Figure 18.**
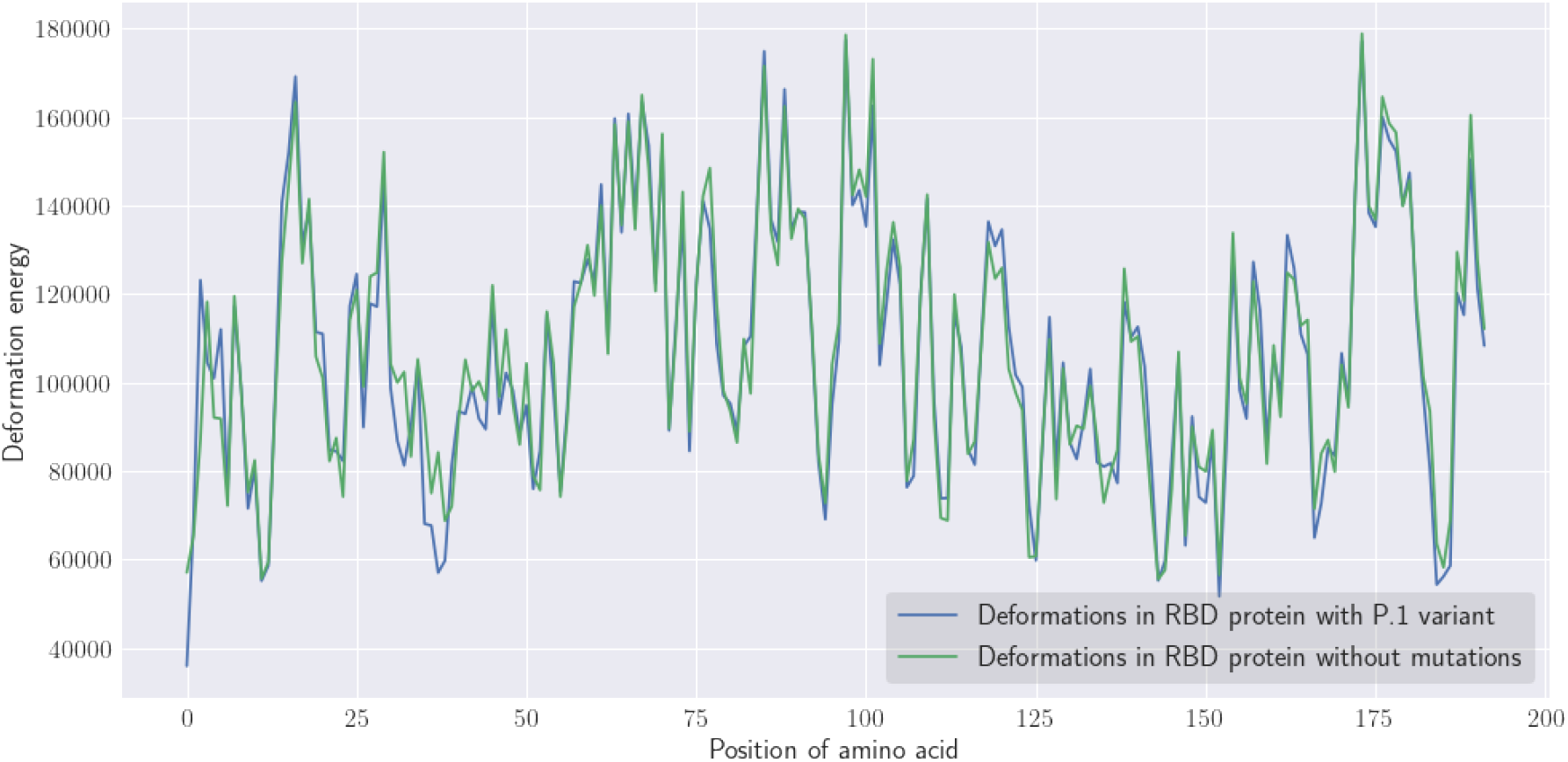
The deformation energies correspond to the last frame of molecular dynamics at 50*ns* only for the RBD region containing the P.1 variant and no mutations. All calculations were performed using the WebNMA tool Tiwari et al. (2014).

#### 3.3.2. Impacts on electrostatic distribution

Based on the physico-chemical properties inherent to each amino acid (Nelson & Cox, 2017), we realized that the transition from the Lys417 residue (*pI ≈* 9.47) positively charged, of a polar character and basic pH to the amino acid Asn417 (*pI ≈* 5.41) also polar although electrically neutral would increase the likelihood of hydrogen bonding at the expense of saline bridges. Similarly, the substitution in line P.1 for Thr417 (*pI ≈* 5.60), and as a consequence of the Threonine being less susceptible to the formation of a saline bridge, there will therefore be a greater probability of accessibility to the solvent as also already discovered at Dejnirattisai et al. (2021). The substitution of the amino acid Asn501 for Tyr501 (*pI ≈* 5.63) apparently would not induce significant differences since both are polar and electrically neutral, therefore having similar probabilities in the formation of hydrogen bonds, except that Tyrosine has an amphipathic character. Finally, the amino acid Glu484 (*pI ≈* 3.08) of a nonpolar character, acidic in addition to being negatively charged when changed by Lys484 with a positive net electrical charge and basic character could therefore alter the electrostatic distribution around it and increase the probability of solubility both in acidic and alkaline conditions where *pI > pH*. From the electrostatic distribution surface (see Figure 19 and Figure 20) calculated by the APBS (Adaptive Poisson-Boltzmann Solver) algorithm (Jurrus et al., 2018) for the last frame, we can see that the P.2 strain contributed to there being regions with a more negative potential (−597.134*kT* to +549.126*kT*) possibly as a result of the E484K mutation. Meanwhile, the behavior of variant P.1 was markedly more negative (−604.138*kT* to +529.056*kT*). On the other hand, the wild-type structure showed a slightly more positive distribution (−618.150*kT* to +535.559*kT*). We believe that this change in electrical charge in some residues may have led to greater electrostatic complementarity in regions that have undergone mutation, which would therefore lead to an increase in affinity in the ACE2-RBD interaction, especially in relation to P.1, something that has already been verified experimentally (Dejnirattisai et al., 2021).

**Figure 19.**
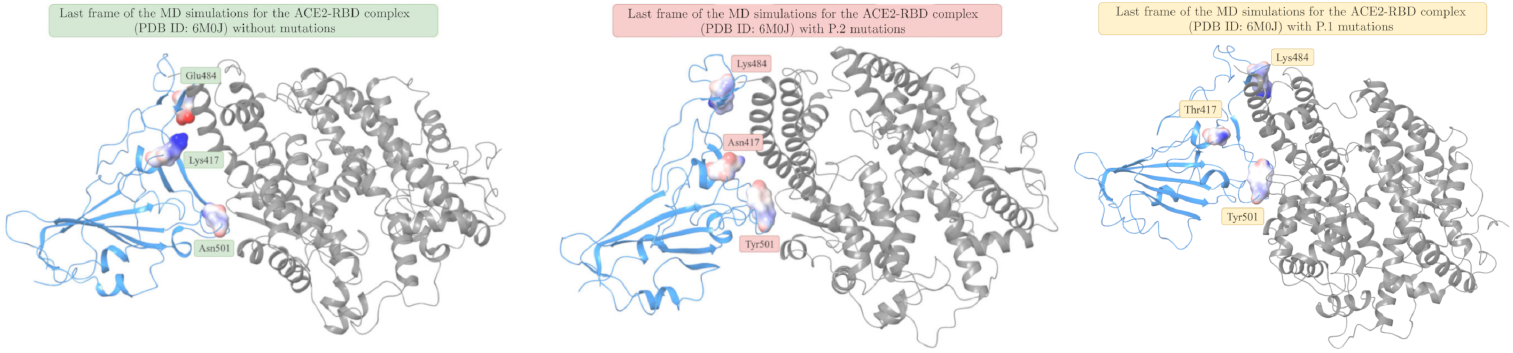
Comparison between the electrostatic distribution surfaces generated with the APBS algorithm from the last frames of the respective molecular dynamics simulations for the ACE2-RBD complex (PDB ID: 6M0J). The visualization was using Schrödinger Maestro 2020-4 software.

**Figure 20.**
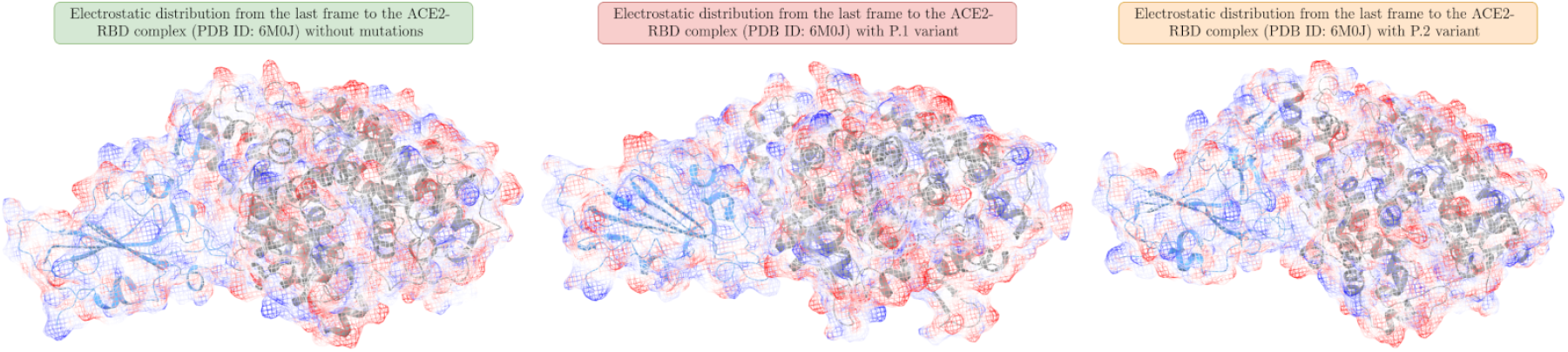
Electrostatic distribution calculated by the APBS algorithm for the entire ACE2-RBD complex (PDB ID: 6M0J). A comparison between the last frames of the P.1, P.2 variant is presented, in addition to the wild-type structure.

In the qualitative aspect, we noticed that the P.1 strain made the *β*-sheet formed between the K417N and N501Y mutations with a shorter length compared to the structure absent from mutations. In addition, there was the disappearance of a *α*-helix between the Ser366 and Asn370 residues, structural factors that may be correlated with the greater transmissibility of the P.1 variant. From the visualization in Figure 19, we realized that the K417T substitution occurred in a loop region that is highly structurally unstable, and therefore, the greater the recurrence of mutations. In addition, we must note that the K417T and E484K mutations are all close to the interface with the ACE2 receptor as analyzed in the PDBePISA Krissinel (2010) platform, which makes them so worrying and more susceptible to the formation of intermolecular interactions.

Finally, it is important to remember that the impacts of strains P.1 and P.2 will become clearer as there is a repetition of the simulations or an increase in the time interval, which would generate more conclusive results, convergence of fluctuations and greater reproducibility. Therefore, among the future perspectives, it is precisely the continuation of simulations to deepen the understanding at the theoretical level of the impacts of the emerging variants of SARS-CoV-2.

## 4. Conclusions

In general, the P.1 variant provided greater stabilization of the ACE2-RBD complex supported by 3 (three) analyzes of molecular dynamics, lower mean RMSF values, greater formation of hydrogen bonds and less exposure to solvent measured by the SASA value. These results reflect less entropic penalty and therefore greater susceptibility to seasonal influences as a result of the term −*T ·* Δ*S* in addition to the possible increase in interaction affinity and greater probability of cell entry. Through the MM-PBSA energy decomposition, we noticed that Van der Waals interactions predominated and were more favorable when the structure has P.1 variant mutations. Consequently, a priori, the variants will take some time to disappear due to evolutionary pressure, as they benefit the virus, and perhaps because of this there is a convergence of certain mutations such as E484K and N501Y that were theoretically explained by molecular dynamics. The P.1 concern variant still remains recognizable by the immune system, and therefore, even if the efficacy of the P2B-2F6 antibody induced by vaccines can be reduced according to molecular dynamics results, there are no conclusions in the literature that would nullify it. In this research, it was noticed a smaller formation of hydrogen bonds in relation to P.1. As long as we don’t completely stop the virus, the possibility of the emergence of new variants is imminent. Therefore, the vaccination of the greatest number of citizens is essential to eradicate the pandemic. Finally, we hope with these results to help in the theoretical understanding of the impacts of emerging lineages in Brazil and also around the world. Therefore, it is essential that there is social distance while the pandemic is not eradicated, in order to minimize the emergence of new mutations specifically in the Spike glycoprotein.

## Acknowledgements

We have sincere gratitude for all the people who came together to face this difficult period of COVID-19. We would like to thank National Council for Scientific and Technological Development (CNPq) for supplying this research with grant 136222/2020-0.

## Supplementary material

All supplementary material for this research can be found at (https://zenodo.org/record/5111905).

## Author contributions statement

*Kelson M. T. Oliveira:* led the project, review and article writing; *Micael D. L. Oliveira:* performed simulations in molecular dynamics/thermodynamics stability with NAMD3, and thus takes responsibility for data integrity and analysis accuracy; *Jonathas N. Silva:* performed molecular dynamics simulations with GROMACS, validated all the results and discussions carried out in the research; *Rosiane de Freitas:* writing-review, editing and development of reconstruction algorithm; *Clarice Santos:* developed the protein reconstruction algorithm. *João Bessa:* Helped scientific discussions throughout the project.

