## Supplementary Material for "Theoretical causes of the Brazilian P.1 and P.2 lineages of the SARS-CoV-2 virus through molecular dynamics" (from another version of this preprint: version 1, 2021-04-10)

[a] Laboratory of Theoretical and Computational Chemistry. Department of Chemistry, Federal University of Amazonas, 69077-000, Manaus, AM, Brazil;

[b] Institute of Computing (IComp), Federal University of Amazonas, 69077-000, Manaus, AM, Brazil.

**ARTICLE HISTORY**

Compiled April 2, 2021

**KEYWORDS**

SARS-CoV-2, molecular dynamics, mutagenesis, P.1/P.2 lineages.

**Table 1.** In order to justify the validity of the Student's t-test approach, the Shapiro-Wilk normality test was performed for the quantity of hydrogen bonds, with a minimum confidence interval of 87%. Levene's test was applied to verify equality between variances where $p > 0.05$, and therefore the inequality had to be considered throughout the hypothesis test. Consequently, we believe that the proposal of the t-test formalism for comparative analysis of MD simulations is justified. Regarding the RMSD and RMSF data set, the 95% confidence interval was adopted.

| | | **p-value** | | | | | |
|---|---|---|---|---|---|---|---|
| Complex | PDB ID | *RMSD* | *RMSF* | Hydrogen Bonds | SASA | $Q(t)$ | *Rg* |
| ACE2-RBD | 6M0J | $p < 0.05$ | $p < 0.05$ | $p < 0.05$ | $p > 0.05$ | $p < 0.05$ | $p < 0.05$ |
| IgG | 7BWJ | $p < 0.05$ | $p < 0.05$ | $p < 0.05$ | - | $p < 0.05$ | $p < 0.05$ |

CONTACT

**Table 2.** The same statistical approaches previously mentioned were used in the comparison of energy data for MM-PBSA decomposition.

| Complex | PDB ID | p-value | | |
|---|---|---|---|---|
| | | $\langle E_{elec} \rangle$ | $\langle E_{VdW} \rangle$ | $\langle E_{MM} \rangle$ |
| ACE2-RBD | 6M0J | $p < 0.05$ | $p < 0.05$ | $p < 0.05$ |
| IgG | 7BWJ | $p < 0.05$ | $p < 0.05$ | $p < 0.05$ |

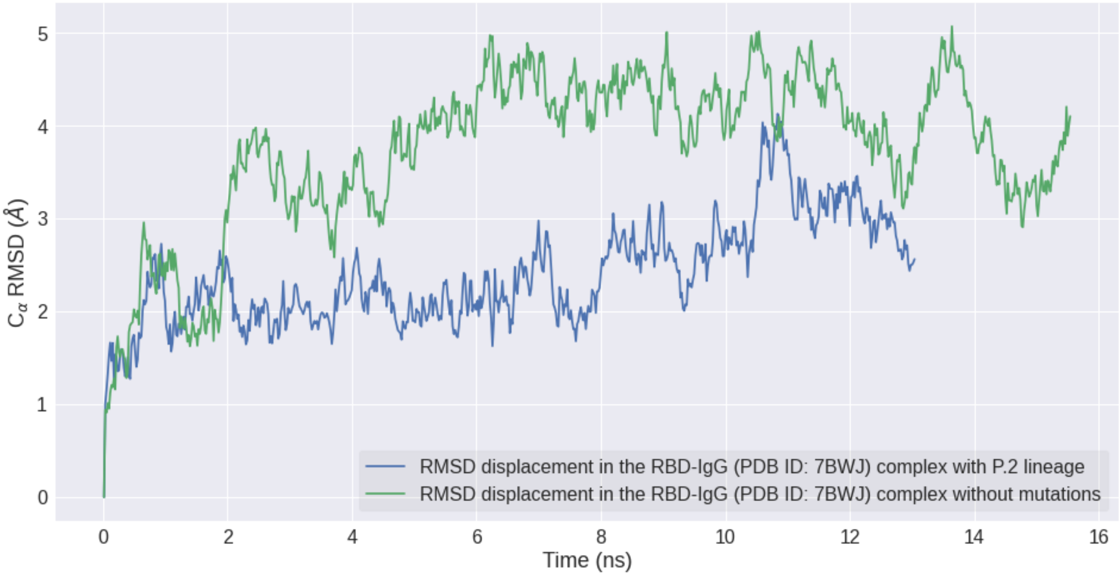


**Figure 1.** Comparison of RMSD values for the RBD-IgG complex (PDB ID: 7BWJ) regarding the P.2 strain.

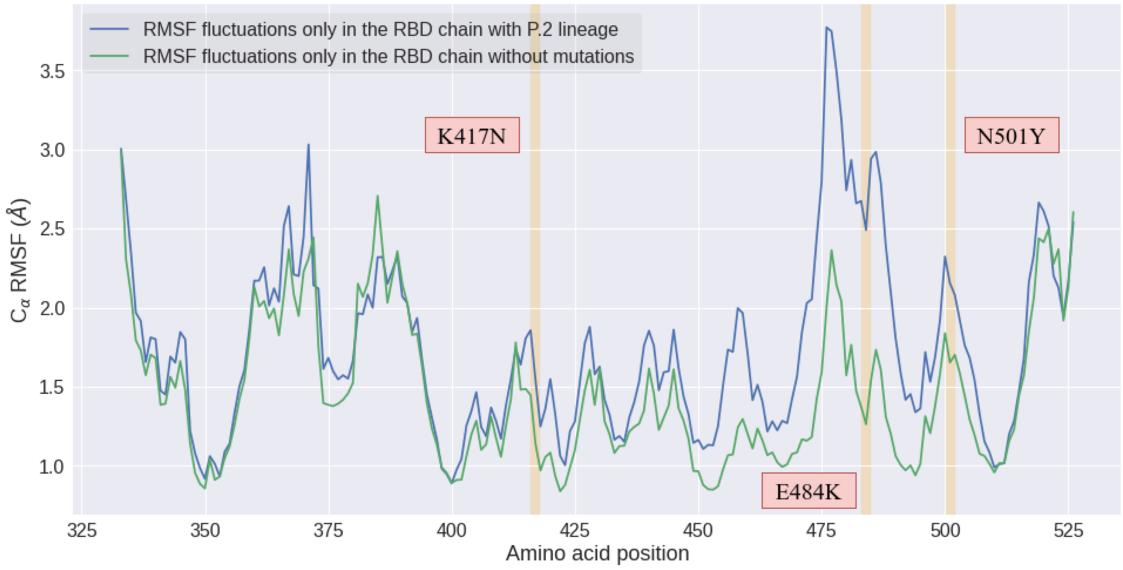


**Figure 2.** Comparison of RMSF values for the RBD-IgG complex (PDB ID: 7BWJ) regarding the P.2 strain.

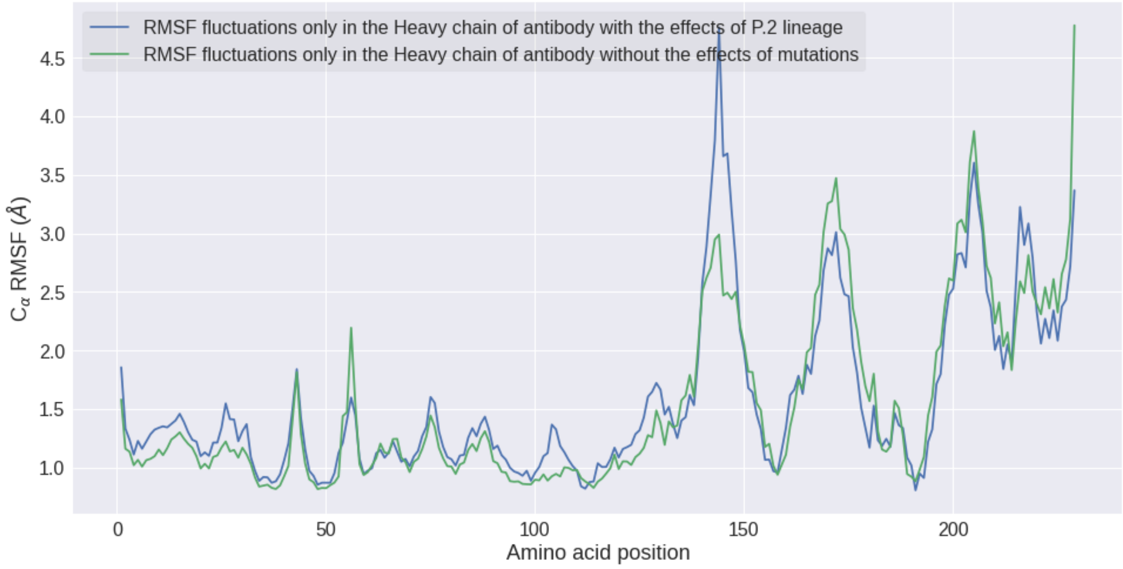


**Figure 3.** Comparison of RMSF values for the complex between the antibody heavy chain and RBD (PDB ID: 7BWJ) regarding the P.2 strain.

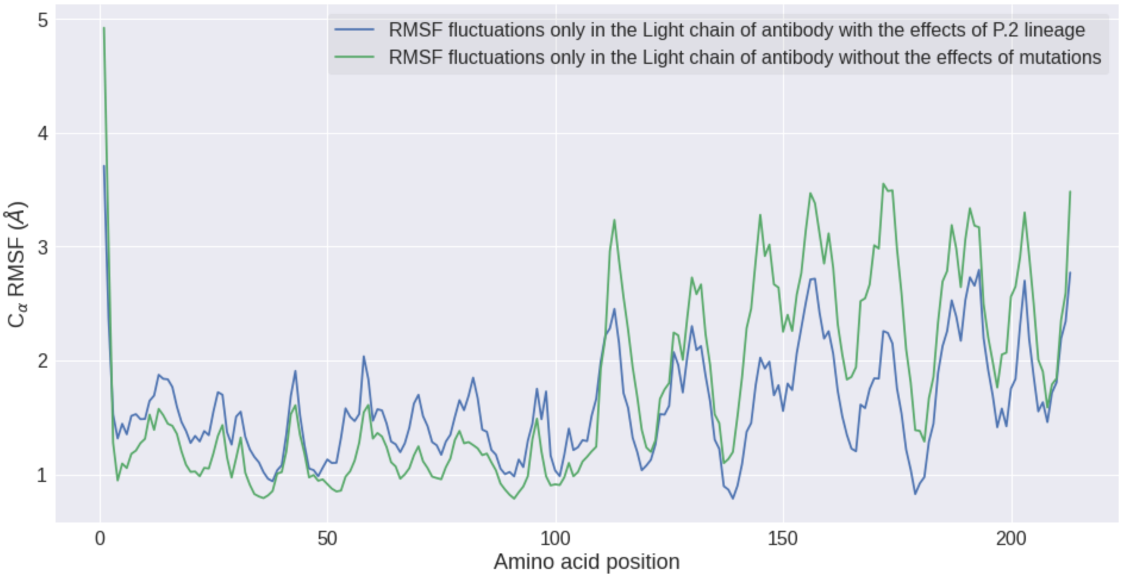


**Figure 4.** Comparison of RMSF values for the complex between the antibody light chain and RBD (PDB ID: 7BWJ) regarding the P.2 strain.

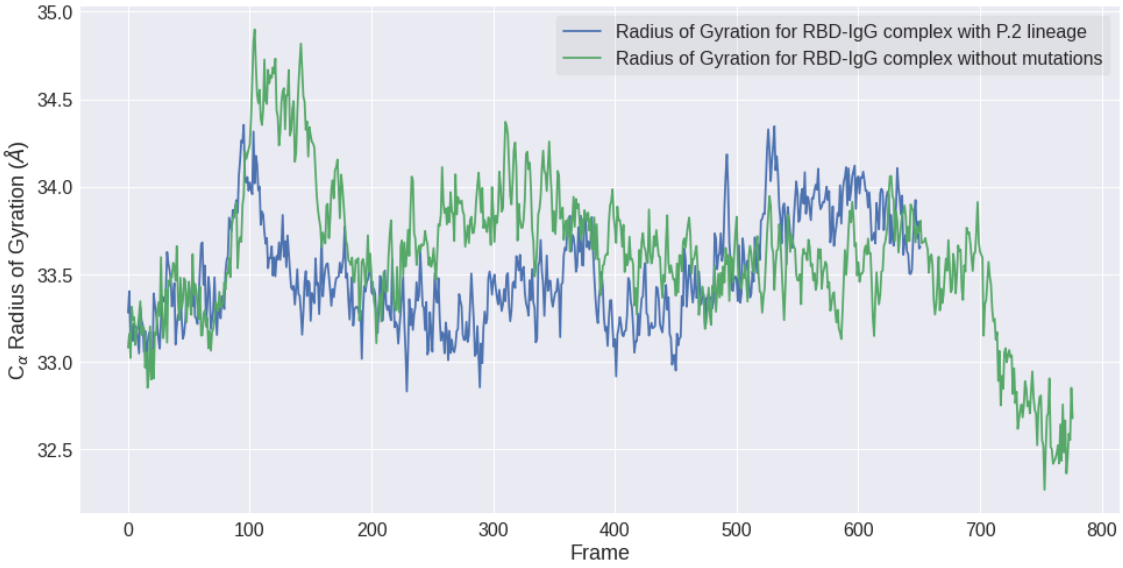


**Figure 5.** Comparison of Radius of Gyration (Rg) values for the RBD-IgG complex (PDB ID: 7BWJ) regarding the P.2 strain.

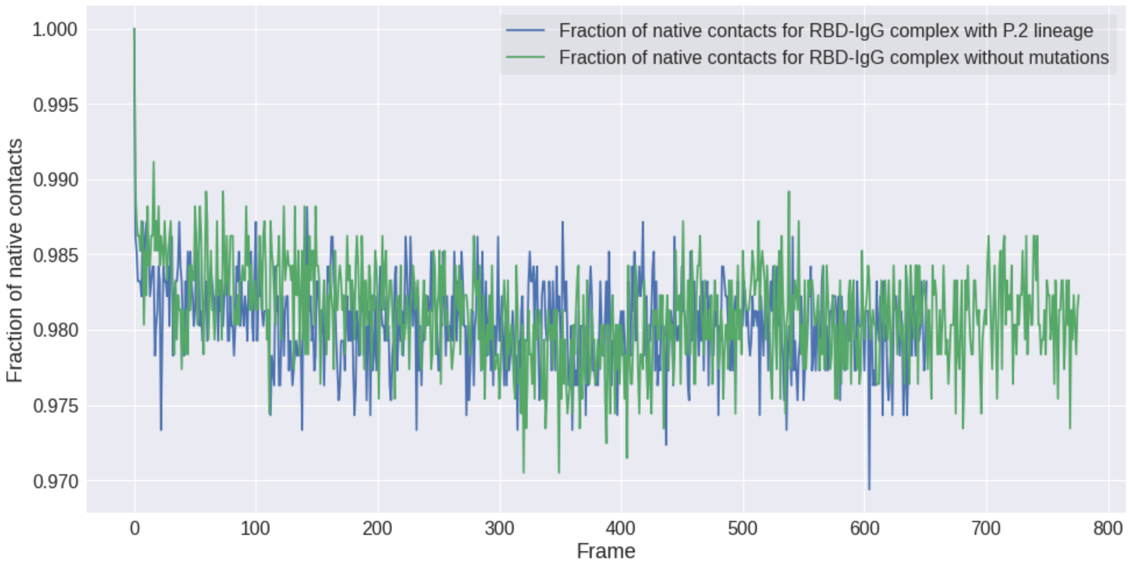


**Figure 6.** Comparison between the values of the fraction of native contacts $Q(t)$ for the RBD-IgG complex (PDB ID: 7BWJ) regarding the P.2 strain.

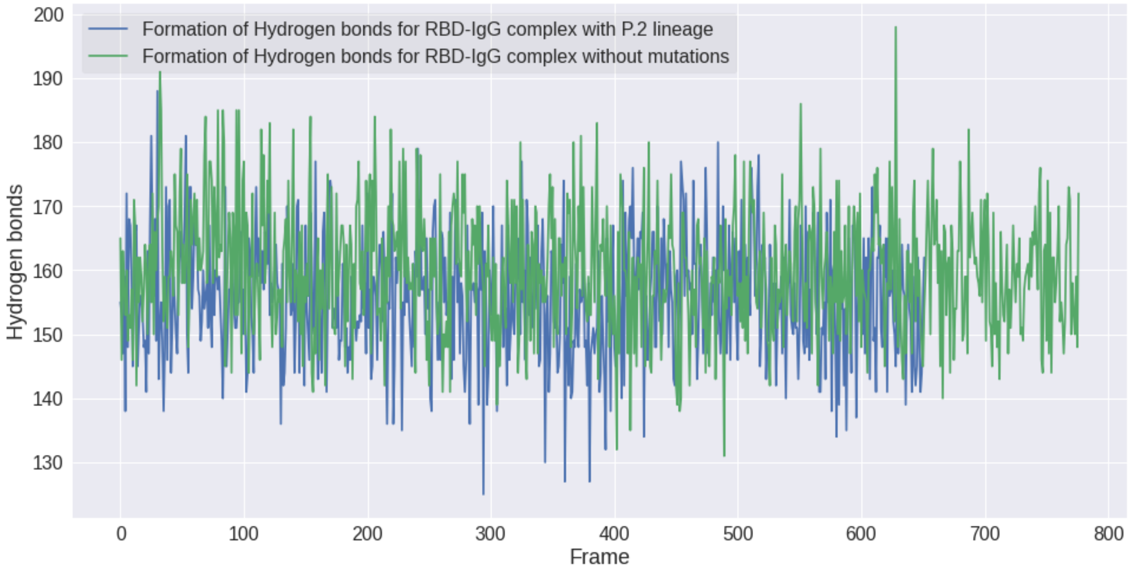


**Figure 7.** Comparative in the formation of Hydrogen bonds for the RBD-IgG complex (PDB ID: 7BWJ) regarding the P.2 strain.

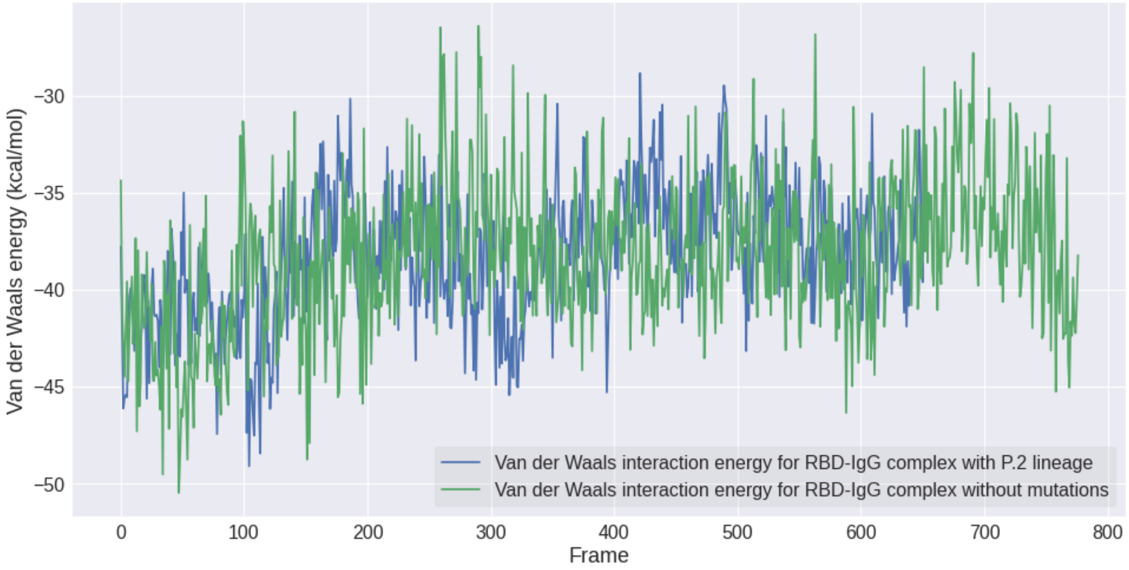


**Figure 8.** Comparison on Van der Waals interaction energy between the antibody heavy chain and RBD (PDB ID: 7BWJ) regarding the P.2 strain.
